# Excitatory and inhibitory subnetworks are equally selective during decision-making and emerge simultaneously during learning

**DOI:** 10.1101/354340

**Authors:** Farzaneh Najafi, Gamaleldin F Elsayed, Robin Cao, Eftychios Pnevmatikakis, Peter E. Latham, John P Cunningham, Anne K Churchland

## Abstract

Inhibitory neurons, which play a critical role in decision-making models, are often simplified as a single pool of non-selective neurons lacking connection specificity. This assumption is supported by observations in primary visual cortex: inhibitory neurons are broadly tuned in vivo, and show non-specific connectivity in slice. Selectivity of excitatory and inhibitory neurons within decision circuits, and hence the validity of decision-making models, is unknown. We simultaneously measured excitatory and inhibitory neurons in posterior parietal cortex of mice judging multisensory stimuli. Surprisingly, excitatory and inhibitory neurons were equally selective for the animal’s choice, both at the single cell and population level. Further, both cell types exhibited similar changes in selectivity and temporal dynamics during learning, paralleling behavioral improvements. These observations, combined with modeling, argue against circuit architectures assuming non-selective inhibitory neurons. Instead, they argue for selective subnetworks of inhibitory and excitatory neurons that are shaped by experience to support expert decision-making.

## Introduction

In many decisions, noisy evidence is accumulated over time to support a categorical choice. At the neural level, there are a number of models that can implement evidence accumulation (Wang, 2002; Machens et al., 2005; Bogacz et al., 2006; Lo and Wang, 2006; Wong and Wang, 2006; Beck et al., 2008; Lim and Goldman, 2013; Rustichini and Padoa-Schioppa, 2015; Mi et al., 2017). Although these circuit models have successfully reproduced key characteristics of behavioral and neural data during perceptual decision-making, their empirical evaluation has been elusive, mainly due to the challenge of identifying inhibitory neurons reliably and in large numbers in behaving animals. Inhibition, which constitutes an essential component of these models, is usually provided by a single pool of inhibitory neurons receiving broad input from all excitatory neurons (non-selective inhibition, Deneve et al., 1999; Wang, 2002; Mi et al., 2017).

The assumption of non-selective inhibition in theoretical models was, perhaps, motivated by some empirical studies that examined the connectivity and tuning of inhibitory and excitatory neurons. Many studies in primary visual cortex report that inhibitory neurons have, on average, broader tuning curves than excitatory neurons for visual stimulus features such as orientation (Sohya et al., 2007; Niell and Stryker, 2008; Liu et al., 2009; Kerlin et al., 2010; Bock et al., 2011; Hofer et al., 2011; Atallah et al., 2012; Chen et al., 2013; Znamenskiy et al., 2018), spatial frequency (Niell and Stryker, 2008; Kerlin et al., 2010; Znamenskiy et al., 2018), and temporal frequency (Znamenskiy et al., 2018). The broad tuning in inhibitory neurons has been mostly attributed to their dense (Hofer et al., 2011; Packer and Yuste, 2011) and functionally unbiased inputs from the surrounding excitatory neurons (Kerlin et al., 2010; Bock et al., 2011; Hofer et al., 2011). This is in contrast to excitatory neurons, which show relatively sharp selectivity to stimulus features (Sohya et al., 2007; Niell and Stryker, 2008; Ch’ng and Reid, 2010; Kerlin et al., 2010; Hofer et al., 2011; Isaacson and Scanziani, 2011; Lee et al., 2016), reflecting their specific and non-random connectivity (Yoshimura et al., 2005; Ch’ng and Reid, 2010; Hofer et al., 2011; Ko et al., 2011; Cossell et al., 2015; Ringach et al., 2016).

Based on the relatively weak tuning of inhibition, it seems reasonable to assume that inhibition in decision circuits is non-specific. However, the overall picture from experimental observations is more nuanced than the original studies would suggest. First, a number of V1 studies report tuning of inhibitory neurons that is on par with excitatory neurons (Ma et al., 2010; Runyan et al., 2010), likely supported by targeted connectivity with excitatory neurons (Yoshimura and Callaway, 2005). Strong tuning of inhibitory neurons has also been reported in primary auditory cortex (Moore and Wehr, 2013). Further, interneurons have been shown to selectively represent key task parameters in behaving animals in areas beyond sensory cortices. In frontal and parietal areas, interneurons can distinguish go vs. no-go responses (For example, Allen et al., 2017) as well as the trial outcome (Pinto and Dan, 2015). Similarly, in the hippocampus, interneurons have strong selectivity for the stimulus (Lowett-Brown 2017), and the animal’s location (Maurer et al., 2006; Ego-Stengel and Wilson, 2007).

This selectivity of inhibitory neurons in a wealth of areas and conditions argue that the assumption of non-selective interneurons in decision-making models must be revisited. Here, we aimed to evaluate this assumption directly. We compared the selectivity of inhibitory and excitatory neurons in PPC of mice during rate discrimination decisions. Surprisingly, we found that excitatory and inhibitory neurons in PPC are equally choice-selective. Moreover, during learning, the specificity of excitatory and inhibitory neurons increased in parallel. These results constrain decision-making models, and in particular argue that in decision areas, subnetworks of selective inhibitory neurons emerge during learning and are engaged during expert decisions.

## Results

To test how excitatory and inhibitory neurons coordinate during decision-making, we measured neural activity in transgenic mice trained to report decisions about the repetition rate of a sequence of multisensory events by licking to a left or right waterspout (Figure 1A; Figure S1A). Trials consisted of simultaneous clicks and flashes, generated randomly (via a Poisson process) at rates that ranged from 5 to 27 Hz over a 1000 ms period (Brunton et al., 2013; Odoemene et al., 2017). Mice reported whether event rates were high or low compared to an abstract category boundary (16 Hz) that they learned from experience. Decisions depended strongly on stimulus rate: performance was at chance when the stimulus rate was at the category boundary, and was better at rates further from the category boundary (Figure 1B). A logistic regression model demonstrated that choice depends on the current stimulus strength, previous choice outcome (Hwang et al., 2017), and the time elapsed since the previous trial (Figure S1B).

**Figure 1.**
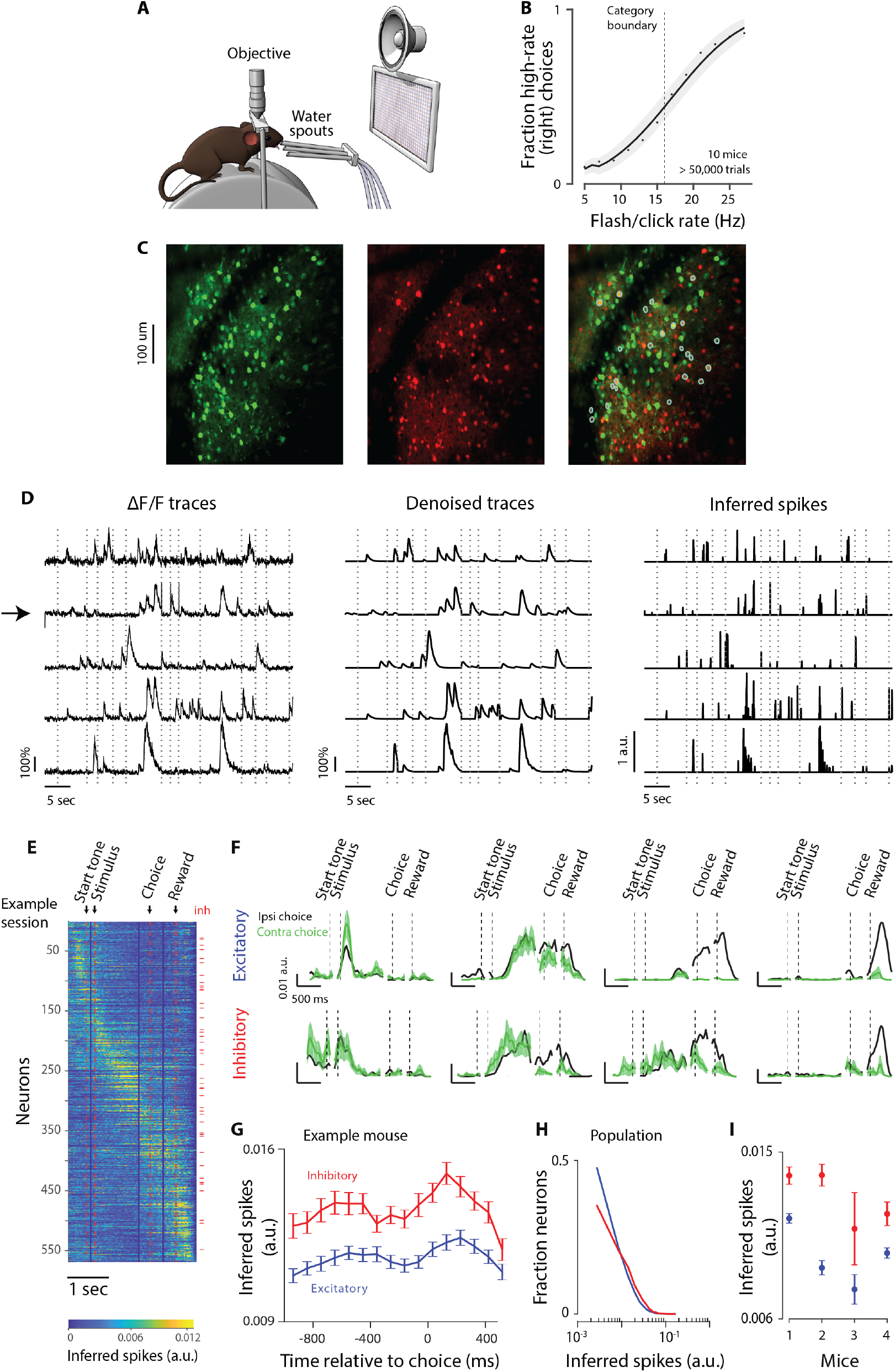
Simultaneous imaging of inhibitory and excitatory populations during decision-making to test decision-making models. **A.** Behavioral apparatus in which a head-fixed mouse is atop a cylindrical wheel. Visual display and speaker present the multisensory stimulus. To initiate a trial, mice licked the middle waterspout. To report the decision about the stimulus rate, mice licked left/right spouts. Objective belongs to the 2-photon microscope used to image neural activity through a window implanted in the skull. **B.** Psychometric function showing the fraction of trials in which the mouse judged the stimulus as high rate as a function of stimulus rate. Dots: data, mean across 10 mice. Line: Logit regression model fit using glmfit.m; mean across mice. Shaded area: standard deviation of the fit across mice. Dashed vertical line: category boundary (16Hz). **C**, Average image of 10,000 frames. **Left:** green channel showing GCaMP6f expression. **Middle:** red channel showing tdTomato expression. **Right:** merge of left and middle. Cyan circles indicate GCaMP6f-expressing neurons that were identified as inhibitory. **D**, Five example neurons identified by the CNMF algorithm (arrow: inhibitory neuron). **Left:** raw ΔF/F traces. **Middle:** de-noised traces. **Right:** inferred spiking activity. Imaging was not performed during inter-trial intervals; traces from 13 consecutive trials are concatenated; dashed lines: trial onsets. **E**, Example session with 568 neurons. Each row shows the trial-averaged inferred spiking activity of a neuron (frame resolution: 32.4ms). Neurons are sorted according to the timing of their peak activity. To ensure peaks were not driven simply by noisy fluctuations, we first computed trial-averaged activity using half of the trials for each neuron. We then identified the time of peak activity for the trial-averaged response. Finally, these peak times were used to determine the plotting order for the trial-averaged activity corresponding to the remaining half of the trials. This crossvalidated approach ensured that the tiling appearance of peak activities was not due to the combination of sorting and false-color-plotting. Inhibitory neurons (*n*=45) are indicated by red ticks on the right. Red vertical lines mark trial events: initiation (start) tone, stimulus onset, choice, and reward. Duration between events (e.g. between start tone and stimulus) varied across trials; so in order to make trial-averaged traces that represent how neural activity changes following trial events (e.g. start tone, stimulus, etc), traces were separately aligned to each trial event, and then averaged across trials. Next, these averaged traces (each aligned to a different trial event) were concatenated to represent neural activity during the entire trial duration, and in response to different trial events. Vertical blue lines indicate the border between the concatenated traces. **F**, Trial-averaged inferred spiking activity of 4 excitatory (top) and 4 inhibitory (bottom) neurons, for ipsi- (black) and contralateral (green) choices (mean +/- standard error; ~250 trials per session). **G**, Inferred spiking activity for excitatory (blue) and inhibitory (red) neurons during the course of a trial. Example mouse; mean +/- standard error across days (*n*=46). Each point corresponds to an average over trials and neurons. Inferred spiking activity was initially downsampled by averaging over three adjacent frames (Methods). Spiking activity was significantly higher for inhibitory neurons (t-test; p<0.001) at all times. **H**, Distribution of inferred spiking activity at time bin 0-97ms (averaged over the three frames before the choice) for all mice and all sessions (41,723 excitatory and 5,142 inhibitory neurons). **I**, Inferred spiking activity at time bin 0-97ms before the choice for each individual mouse (mean +/- standard error across days). Differences were significant for all subjects (t-test; p<0.001).

We imaged excitatory and inhibitory neural activity by injecting a viral vector containing the calcium indicator GCaMP6f to layer 2/3 of mouse Posterior Parietal Cortex (PPC; 2mm posterior to Bregma, 1.7mm lateral to midline (Harvey et al., 2012; Funamizu et al., 2016; Goard et al., 2016; Morcos and Harvey, 2016; Hwang et al., 2017; Song et al., 2017)). Mice expressed the red fluorescent protein tdTomato transgenically in all GABAergic inhibitory neurons. We used a two-channel two-photon microscope to record the activity of all neurons, a subset of which were identified as inhibitory neurons (Figure 1C). This allowed us to measure the activity of excitatory and inhibitory populations in the same animal.

To detect neurons and extract calcium signals from imaging data, we leveraged an algorithm that simultaneously identifies neurons, de-noises the fluorescence signal and de-mixes signals from spatially overlapping components (Pnevmatikakis et al., 2016; Giovannucci et al., 2018) (Figure 1D middle). The algorithm also estimates spiking activity for each neuron, yielding, for each frame, a number that is related to the spiking activity during that frame (Figure 1D right). We refer to this number as “inferred spiking activity”, acknowledging that estimating spikes from calcium signals is challenging (Chen et al., 2013). In particular, while higher inferred spiking activity within a single neuron indicates higher firing rates, comparison of firing rates across neurons is not possible with this method. Analyses were performed on inferred spiking activity. To identify inhibitory neurons, we used a method that we developed to correct for bleed-through from the green to the red channel (Methods). Next, we identified a subset of GCaMP6f-expressing neurons as inhibitory neurons based on the signal intensity on the red channel as well as the spatial correlation between red and green channels (Figure 1C right, cyan circles). Inhibitory neurons constituted 11% of the population, within the range of the previous reports (Beaulieu, 1993; Gabbott et al., 1997; Rudy et al., 2011; Sahara et al., 2012), but on the lower side due to our desire to be conservative in assigning neurons to the inhibitory pool (Methods).

Confirming previous reports (Funamizu et al., 2016; Morcos and Harvey, 2016; Runyan et al., 2017), we observed that the activity of individual neurons peaked at time points that spanned the trial (Figure 1E,F). Diverse temporal dynamics were evident in both cell types (Figure 1E,F) and did not appreciably differ between the two (Figure S2). The magnitude of inferred spiking activity was significantly different for inhibitory compared to excitatory neurons throughout the trial (Figure 1G; t-test, p<0.001). In the moments before the choice (97.1ms, average of 3 frames), this difference was clear (Figure 1H) and significant for all mice (Figure 1I). The probable differences in GCaMP expression levels and calcium buffering between excitatory and inhibitory neurons, as well as how spiking activity is inferred (Methods), precludes a direct estimate of the underlying firing rates (Kwan and Dan, 2012). However, the significant difference in the inferred spiking activity between excitatory and inhibitory neurons provides further evidence that we successfully identified two separate neural populations.

### Individual excitatory and inhibitory neurons are similarly choice-selective

To assess the selectivity of individual excitatory and inhibitory neurons for the decision outcome, we performed receiver operating characteristic (ROC) analysis (Green and Swets, 1966) on single-neuron responses. For each neuron, at each time point, we calculated the area under the ROC curve (AUC) as a measure of the amount of overlap between the response distributions for ipsilateral vs. contralateral choices. A neuron was identified as “choice-selective” if its AUC value was significantly different (p<0.05) from a constructed shuffled distribution (Figure S3A; Methods), indicating that the neural activity was significantly different for ipsi- vs. contralateral choices (Figure 2A, shaded areas mark choice-selective neurons).

**Figure 2.**
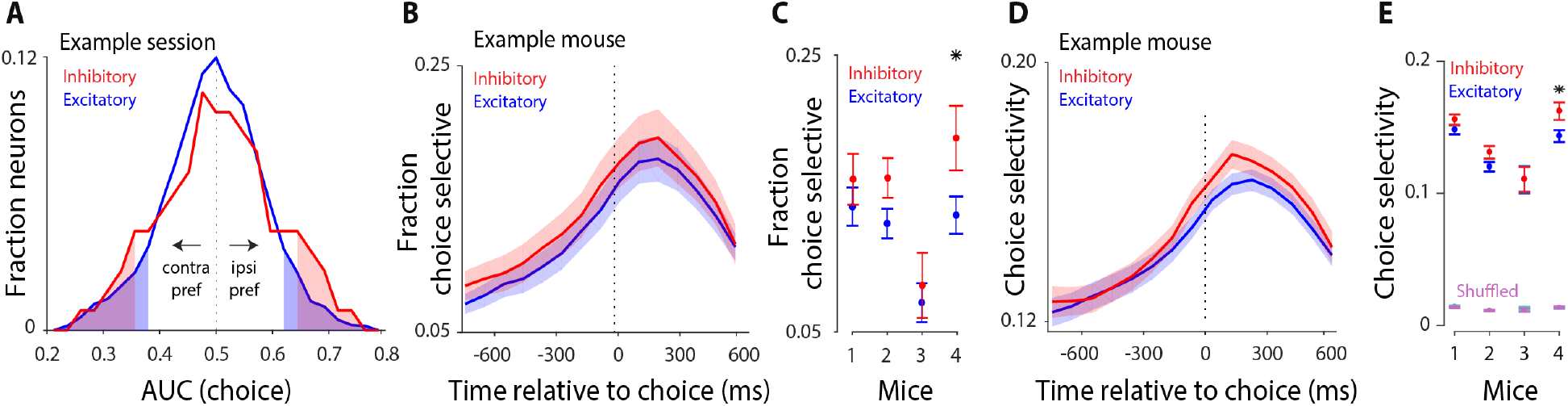
Single-cell and pairwise analyses argue for non-random connections between excitatory and inhibitory neurons. Ideal observer analysis reveals the ability of individual neurons to distinguish left vs. right choices. In all panels, blue and red indicate excitatory and inhibitory neurons, respectively. **A**, Distribution of AUC values (area under the curve) of an ROC analysis for distinguishing choice from the activity of single neurons in an example session. Data correspond to the 97 ms window preceding the choice for 285 excitatory and 29 inhibitory neurons. Values larger than 0.5 indicate neurons preferring the ipsi-lateral choice; values smaller than 0.5 indicate neurons preferring the contralateral choice. Shaded areas mark significant AUC values (compared to a shuffle distribution). Distributions were smoothed (moving average, span=5). For this example session, 5 inhibitory and 24 excitatory neurons were significantly choice selective. **B**, ROC analysis performed on 97 ms non-overlapping time windows. Vertical axis: fraction of excitatory and inhibitory neurons with significant choice selectivity at the corresponding time on the horizontal axis; example mouse; mean+/-standard error across days (n = 45). **C**, Fraction of excitatory and inhibitory neurons that are significantly choice-selective at 0-97 ms before the choice is summarized for each mouse; mean+/-standard error across days (n = 45, 48, 7, 35 sessions per mouse). Star (*) indicates significant difference between excitatory and inhibitory neurons (t-test; p<0.05); see also Figure S3D. Fraction selective neurons at 0-97ms before choice (median across mice): excitatory: 13%; inhibitory: 16%, resulting in ~6 inhibitory and 43 excitatory neurons with significant choice selectivity per session. See also Figure S3C for a different quantification. **D**, ROC analysis performed on 97 ms non-overlapping time windows. Time course of normalized choice selectivity (defined as twice the absolute deviation of AUC from chance) shown for excitatory and inhibitory neurons in an example mouse; mean+/-standard error across days, n=45 sessions. **E**, Average of normalized choice selectivity for excitatory and inhibitory neurons from 0-97 ms before the choice is summarized for each mouse; mean+/-standard error across days. “Shuffled” denotes AUC was computed using shuffled trial labels.

The fraction of choice-selective neurons (Figure 2B) and the magnitude of choice selectivity (Figure 2D) gradually increased during the course of the trial, peaking just after the animal reported its choice. Importantly, excitatory and inhibitory neurons were similar in terms of the fraction of choice-selective neurons (Figure 2B,C; Fig S3B,C), as well as the magnitude and time course (Figure 2D,E) of choice selectivity. These results were not due to differences in inferred spike rates of the two cell types (Figure 1G): when we restricted the ROC analysis to excitatory and inhibitory neurons with similar spiking activity, both cell types remained equally selective for the animal’s choice (Figure S3D).

Next, we assessed whether neurons reflected the animal’s choice or the sensory stimulus, by comparing choice selectivity values resulting from ROC analysis performed on correct vs. error trials. For the majority of neurons, choice selectivity computed on correct trials was similar to that of error trials, resulting in a positive correlation of the two quantities across neurons (Figure S3E). Positive correlations indicate that most neurons reflect the impending choice more so than the sensory stimulus that informed it (Methods). Variability across mice in the strength of this correlation may indicate that the balance of sensory vs. choice signals within individual neurons varied across subjects (perhaps due to imaged subregions within the window, Figure S3E right). Importantly, however, within each subject, this correlation was very similar for excitatory vs. inhibitory neurons (Figure S3E), suggesting that within each animal, the tendency for neurons to be modulated by the choice vs. the stimulus was similar in excitatory and inhibitory neurons.

The existence of task-modulated inhibitory neurons has been reported elsewhere (Maurer et al., 2006; Ego-Stengel and Wilson, 2007; Lovett-Barron et al., 2014; Pinto and Dan, 2015; Allen et al., 2017; Kamigaki and Dan, 2017), but importantly, here choice selectivity was similarly strong in excitatory and inhibitory neurons, both in fraction and magnitude. This was at odds with the commonly accepted assumption of non-specific inhibition in theoretical studies (Deneve et al., 1999; Wang, 2002; Mi et al., 2017), and surprising given the numerous empirical findings, which suggest broad tuning and weakly specific connectivity in inhibitory neurons (Sohya et al., 2007; Niell and Stryker, 2008; Liu et al., 2009; Kerlin et al., 2010; Bock et al., 2011; Hofer et al., 2011; Isaacson and Scanziani, 2011; Packer and Yuste, 2011; Atallah et al., 2012; Chen et al., 2013). This observation was a first hint that specific functional subnetworks, preferring either ipsi- or contralateral choices, exist within the inhibitory population, just like the excitatory population (Yoshimura and Callaway, 2005; Znamenskiy et al., 2018).

### Choice can be decoded with equal accuracy from both excitatory and inhibitory populations

While individual inhibitory neurons could distinguish the animal’s choice about as well as excitatory ones, the overall choice selectivity in single neurons was small (Figure 2E). To further evaluate the discrimination ability of inhibitory neurons, we leveraged our ability to measure hundreds of neurons simultaneously. Specifically, we examined the ability of a linear classifier (support vector machine, SVM; Hofmann et al., 2008) to predict the animal’s choice from the single-trial population activity (cross-validated; L2 penalty; see Methods).

We first performed this analysis on all neurons imaged simultaneously in a single session (Figure 3A, left), training the classifier separately for every moment in the trial (97 ms bins). Classification accuracy gradually grew after stimulus onset and peaked at the time of the choice (Figure 3B, black). Performance was at chance on a shuffle control in which trials were randomly assigned as left or right choice (Figure 3B, shuffled). The ability of the entire population of PPC neurons to predict the animal’s upcoming choice confirms previous observations (Funamizu et al., 2016; Goard et al., 2016; Morcos and Harvey, 2016; Driscoll et al., 2017). Our overall classification accuracy was in the same range as these studies.

**Figure 3.**
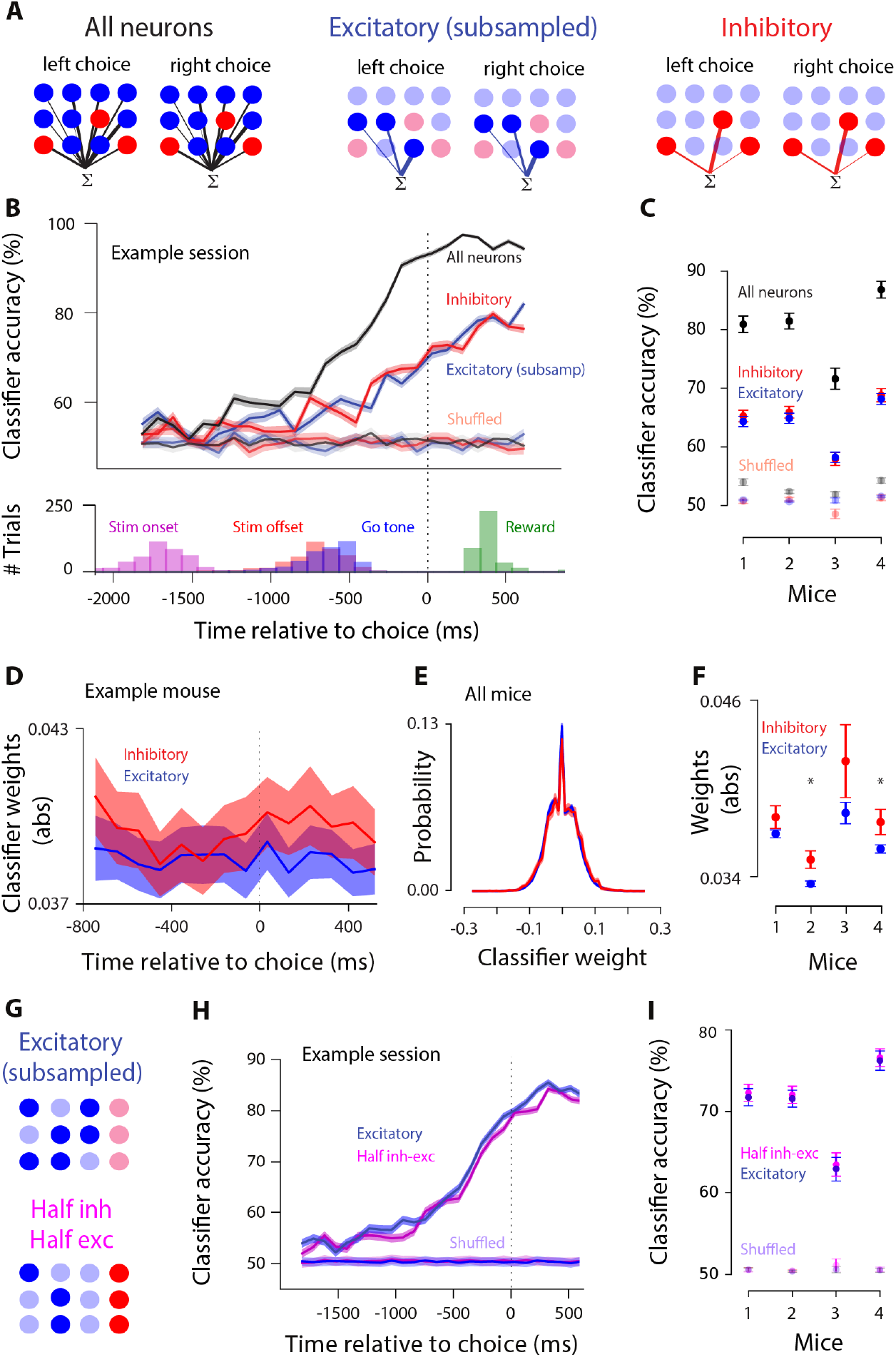
Linear classifiers can predict the animal’s choice with equally high accuracy from the activity of either excitatory or inhibitory populations. **A**, Schematic of decoding choice from the population activity of all neurons (left), only excitatory neurons (middle), subsampled to the same number as inhibitory neurons, and only inhibitory neurons (right). A linear SVM assigns weights of different magnitude (indicated by lines of different thickness) to each neuron in the population so that a weighted sum of population activity differs for trials preceding left vs. right choices. **B, Top:** classification accuracy of decoders trained on all neurons (black), subsampled excitatory neurons (blue), and inhibitory neurons (red) (cross-validated; decoders trained on every 97ms time bin; example session; mean+/-standard error across 50 cross-validated samples). Data are aligned to the animal’s choice (black dotted line). Classification accuracy is lower for inhibitory or subsampled excitatory populations (red, blue) relative to all neurons (black) because of the smaller population size. Classifier accuracy was similar for excitatory and inhibitory populations throughout the trial. Unsaturated lines show performance on shuffled trial labels. **Bottom:** distribution of stimulus onset, stimulus offset, go tone, and reward occurrence for the example session shown on the top. **C**, Classification accuracy during 0-97 ms before the choice for 4 animals on real (saturated) and shuffled (unsaturated) data. Mean+/-standard error across days per mouse. **D**-**F**, When all neurons were included in the decoder (panel A, left), excitatory and inhibitory neurons were assigned weights of similar magnitude. **D**, Absolute value of weights for excitatory and inhibitory neurons in the decoders trained on all neurons, at every moment in the trial; example mouse; mean+/-standard error across days. **E**, Distribution of classifier weights (decoder training time: 0-97 ms before the choice) are similar for excitatory and inhibitory neurons. Neurons from all mice pooled (42,019 excitatory and 5,172 inhibitory neurons). Shading reflects the standard error in each bin of the distribution. **F**, Absolute value of weights in the classifier trained from 0-97 ms before the choice for excitatory vs. inhibitory neurons, for each mouse. Mean+/-standard error across days. Star indicates P<0.05, t-test. **G**, Schematic of decoding choice from a population of subsampled excitatory neurons (top) vs. a population of the same size but including half inhibitory and half excitatory neurons (bottom). **H**, Classifier accuracy of populations including only excitatory (blue) or half inhibitory, half excitatory neurons (magenta); example session. Classifier trained at each moment in the trial. Traces show mean+/-standard error across 50 cross-validated samples. **I**, Summary of each mouse (mean+/-standard error across days) for the decoders trained from 0-97 ms before the choice.

We then examined classifier accuracy for excitatory and inhibitory populations separately. For excitatory neurons, we subsampled the population so that the total number of neurons matched the number of inhibitory neurons in the same session (Figure 3A, middle). As expected, overall classification accuracy was reduced due to the smaller population size; although performance was still well above chance and the temporal dynamics were the same as when all neurons were included (Figure 3B, blue trace). Finally, we included all inhibitory neurons (Figure 3A, right). Surprisingly, the classification accuracy of inhibitory neurons was not only well above chance, but, moreover, was very similar to that of excitatory neurons (Figure 3B, red and blue traces overlap; Figure S4: additional example sessions). Similar classification accuracy for excitatory and inhibitory populations was observed in all subjects (Figure 3C). This result was not due to using inferred spikes: excitatory and inhibitory populations were equally choice selective even when the decoding analysis was performed on calcium traces (Figure S5).

Our analysis may have obscured a difference between excitatory and inhibitory neurons because it evaluated their performance separately, rather than considering how these neurons are leveraged collectively in a classifier that can take advantage of both cell types. To test this, we examined the classifier that was trained on all neurons (Figure 3A left; Figure 3B black), and compared the classifier weights assigned to excitatory vs. inhibitory neurons. We found that the weight magnitudes of excitatory and inhibitory neurons were matched for the entire course of the trial (Figure 3D). Also the distributions of weights were overlapping (Figure 3E,F). The comparable classifier weights for excitatory and inhibitory neurons demonstrate that both cell types were similarly informative about the animal’s upcoming choice.

We next tested whether excitatory and inhibitory populations can be decoded more accurately from a mixed population. This could occur, for example, if the excitatory-inhibitory correlations were weak relative to excitatory-excitatory and inhibitory-inhibitory correlations (Panzeri et al., 1999; Averbeck et al., 2006; Moreno-Bote et al., 2014). To assess this, we trained the classifier on a population that included half excitatory and half inhibitory neurons (Figure 3G bottom), and compared its choice-prediction accuracy with the classifier that was trained on a population of the same size, but consisted only of excitatory neurons (Figure 3G top). We found similar classification accuracy for both decoders during the entire trial (Figure 3H,I), arguing that a mixed population offers no major advantage to decoding.

We next trained new classifiers to evaluate whether population activity reflected additional task features. First, the population activity was somewhat informative about previous trial choice (Figure S6A), in agreement with previous studies (Morcos and Harvey, 2016; Hwang et al., 2017; Akrami et al., 2018); but also see (Zhong et al., 2018). Excitatory and inhibitory populations were similarly selective for the animal’s previous choice (Figure S6A). Second, selectivity for the stimulus category (high rate vs. low rate) was low (Figure S6B), confirming our analysis of correct vs. incorrect trials (single neurons: Figure S3E; population: Figure S6D,E). Again, excitatory and inhibitory populations were similarly selective (Figure S6B). Finally, PPC population activity was strongly selective for the outcome of the trial (reward vs. lack of reward; Figure S6C). Excitatory and inhibitory neurons showed a small but consistent difference in the classifier accuracy (Figure S6C), indicating that once the reward is delivered, the network is operating in a different regime compared to during decision formation, perhaps due to distinct reward-related inputs to excitatory and inhibitory neurons (Pinto and Dan, 2015; Allen et al., 2017). This finding is broadly in keeping with previous studies which suggest that neural populations explore different dimensions over the course of a trial (Raposo et al., 2014; Elsayed et al., 2016).

Finally, we studied the temporal dynamics of the choice signal in PPC population during the course of the trial. If excitatory and inhibitory neurons are connected within subnetworks with frequent cross talk, the two populations should not only predict the animal’s choice with similar accuracy, as shown above, but the readout weights (the weights assigned by the classifier) should exhibit similar temporal dynamics. To assess this, we quantified each population’s stability: the extent to which a classifier trained at one moment could successfully classify neural activity as preceding left vs. right choice at different moments. If population-wide patterns of activity are similar over time (e.g., all neurons gradually increase their firing rates), classifiers trained at one moment will accurately classify neural activity at different moments. Excitatory and inhibitory populations might differ in this regard, with one population more stable than the other.

As the gap between testing and training time increased, a gradual drop occurred in the classifier accuracy, as expected (Figure 4A,B). This drop in accuracy occurred at a very similar rate for excitatory and inhibitory populations (Figure 4B). To quantify this, we determined the time window over which the classifier accuracy remained within 2 standard deviations of the accuracy at the training window (Figure 4C). This was indistinguishable for excitatory and inhibitory neurons (Figure 4D; Figure S7A). An alternate method for assessing stability, computing the angle between the weights of pairs of classifiers trained at different time windows, likewise suggested that excitatory and inhibitory populations are similarly stable (Methods; Figure S7C).

**Figure 4.**
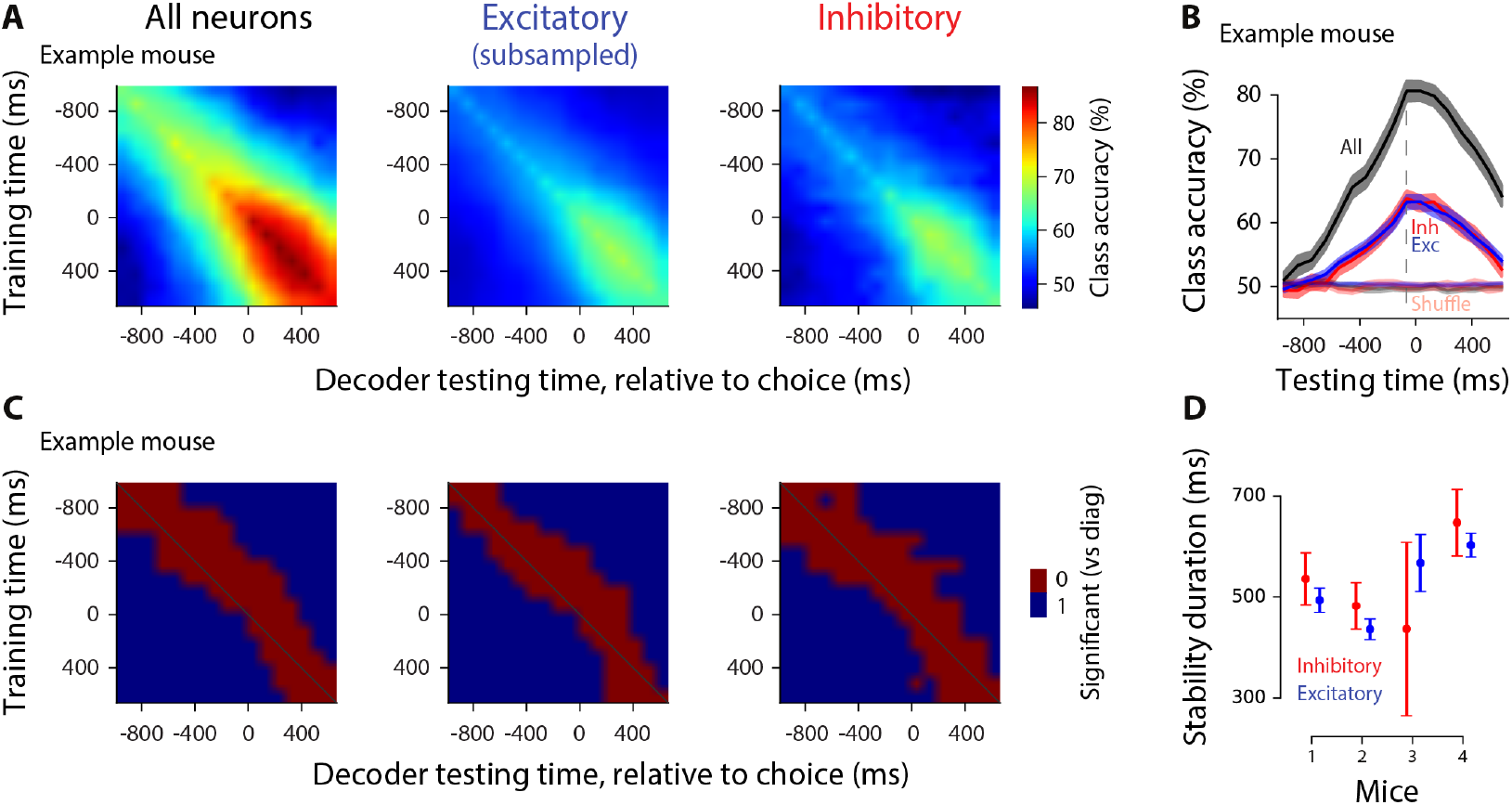
Classifiers, whether trained on excitatory or inhibitory neurons, show comparable stability during decision formation. Cross-temporal generalization of choice decoders. **A**, Classification accuracy of decoders for each pair of training and testing time points, using the population activity of all neurons (left), subsampled excitatory neurons (middle), or inhibitory neurons (right). Diagonal: same training, testing time (same as in Figure 3). Example mouse, mean across 45 sessions. **B**, Example classification accuracy traces showing how classifiers trained at 0-97 ms before choice generalize to other times in the trial. Excitatory and inhibitory neurons show the same time course of generalization. Same mouse as in (A), mean+/-standard error across days **C**, Decoders are stable in a short window outside their training time. Red indicates stability: classification accuracy of a decoder tested at a time different from its training time is within 2 standard deviations of the decoder tested at the same time as the training time. Example mouse; mean across days. **D**, Summary of stability duration for the decoder trained from 0-97 ms before the choice, using inhibitory neurons (red) or subsampled excitatory neurons (blue), for each mouse. Mean+/-standard error across days, per mouse.

### Modeling rules out decision circuits with non-selective inhibition

These results would seem to rule out circuitry from traditional decision-making models, in which the inhibitory neurons are non-selective. This is because in non-selective circuits the average input to the inhibitory neurons is the same whether the evidence favors choice 1 or choice 2 (see Figure 5A, top). However, while the average input is the same, there are fluctuations in connection strength, which can lead to selectivity in some inhibitory neurons. For instance, suppose that, because of the inherent randomness in neural circuits, an inhibitory neuron received more connections from the excitatory neurons in population E_1_ than those in population E_2_. In that case, the firing rate of the inhibitory neuron would be slightly higher when evidence in favor of choice 1 is present. That difference in firing rate could potentially be exploited by a classifier to predict the choice of the animal. Hence, one may argue that even a decision circuit with non-selective inhibition (Figure 5A, top) can lead to similar decoding accuracy in inhibitory and excitatory neurons, questioning whether our experimental findings (Figures 2,3) can be leveraged to constrain decision-making models.

**Figure 5.**
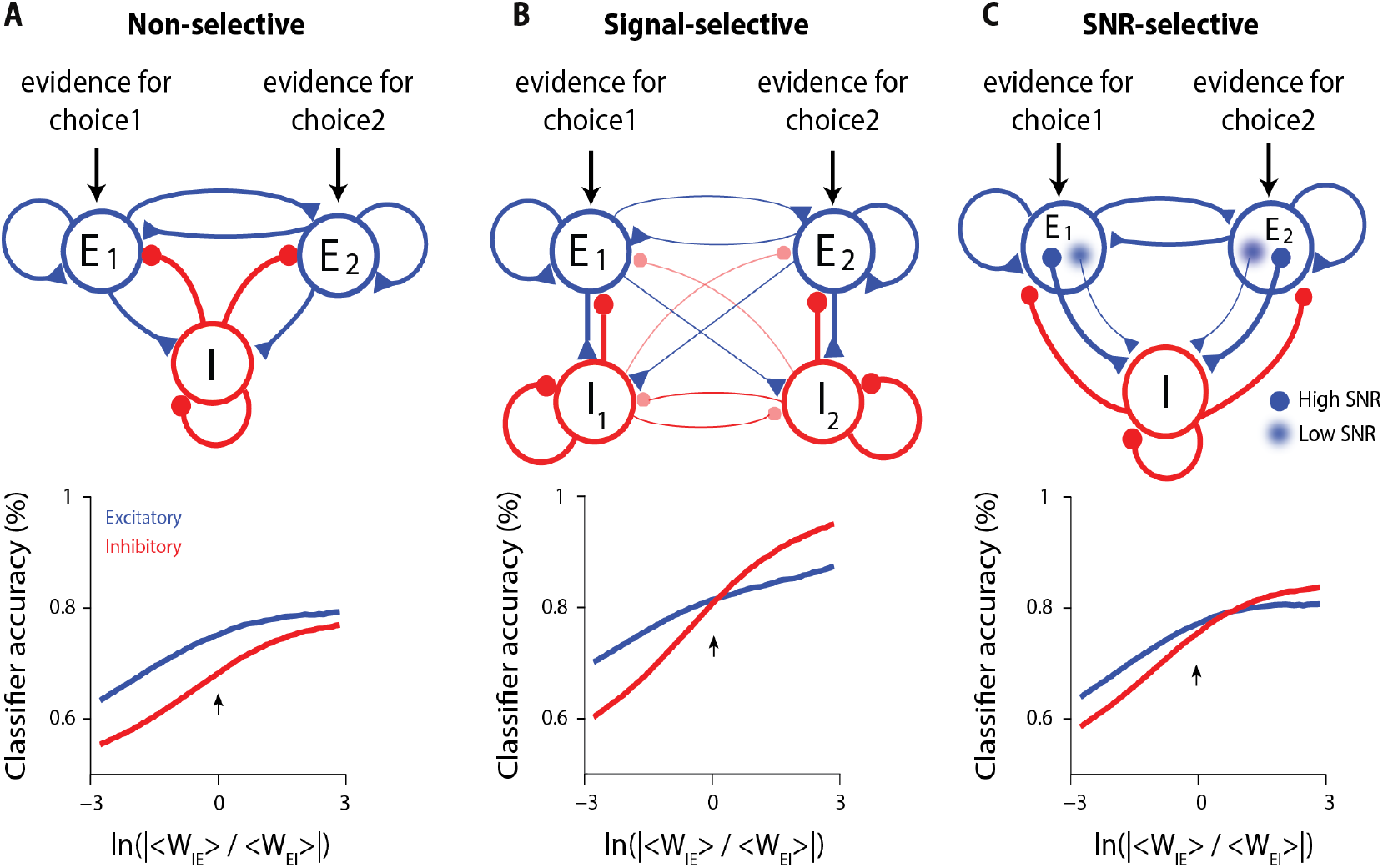
Modeling decision circuits with different architectures. **A, Top**: Non-selective decision-making model. E_1_ and E_2_ represent pools of excitatory neurons, each favoring a different choice. Both pools excite a single pool of non-selective inhibitory neurons (I), which, in turn, provides inhibition to both excitatory pools. **Bottom**: Classification accuracy of excitatory (blue) and inhibitory (red) neurons as a function of the relative strength of excitatory-to-inhibitory vs. inhibitory-to-excitatory connections. For all values of this parameter, excitatory neurons had higher classification accuracy than inhibitory ones. This was true for all parameters tested (Methods; Figure S8; angle brackets denote averages over weights). The arrow in this and subsequent panels indicates the parameter value that is in line with experimental data, which suggest similar connectivity strength for E-to-I and I-to-E connections. **B, Top:** Selective decision-making model. I_1_ and I_2_ represent pools of inhibitory neurons that connect more strongly to E_1_ and E_2_, respectively, than to E_2_ and E_1_, and all cross-pool connections are weaker than within-pool connections. **Bottom**: Decoding accuracy of inhibitory and excitatory neurons match at the biologically plausible regime (arrow). Cross-pool connectivity was 25% smaller than within-pool connectivity. **C, Top**: Selective decision-making model, except now inhibitory neurons connect more strongly to excitatory neurons with high signal to noise ratios (i.e. high input selectivity). **Bottom:** Decoding accuracy of inhibitory and excitatory neurons could match near the biologically plausible regime (arrow). In all panels, decoding accuracy depends on the relative strength of excitatory to inhibitory versus inhibitory to excitatory connections. In (B) and (C), larger excitatory to inhibitory connections favor inhibitory neurons. For all plots we used 50 excitatory and 50 inhibitory neurons out of a population containing 4000 excitatory and 1000 inhibitory neurons.

To test this quantitatively, we modeled a non-selective circuit to evaluate the selectivity of inhibitory neurons in such a circuit architecture (Methods). Classification accuracy depended on the connection strengths between excitatory and inhibitory neurons (horizontal axis on Figure 5A, bottom). This is expected, because large changes in connection strength values can have a large impact on how the network operates. The most biologically plausible regime is near 0, corresponding to the equal strengths for excitatory-to-inhibitory and inhibitory-to-excitatory connections (Thomson and Lamy, 2007; Jouhanneau et al., 2015; Jouhanneau et al., 2018; Znamenskiy et al., 2018) (Figure 5A, arrow). For this value (and indeed for all other values), inhibitory neurons had lower classification accuracy than excitatory neurons (Figure 5A, bottom; Figure S8, left), inconsistent with our experimental results (Figure 3B,C). Therefore, in the non-selective circuit, although some inhibitory neurons can become selective due to random biased inputs from the excitatory pools, the classification accuracy of inhibitory neurons will still be lower than excitatory neurons, regardless of the model parameters. This is because even modest amounts of noise in the system are sufficient to overcome any informative randomness in excitatory to inhibitory connections.

Next, we modeled a signal-selective circuit; in which inhibitory neurons were connected preferentially to one excitatory pool over the other. As a result, selective pools of inhibitory neurons were generated, just like excitatory neurons (Figure 5B, top). In this circuit architecture, inhibitory and excitatory neurons had matched classification accuracy when the connection strength between excitatory and inhibitory neurons was in the biologically plausible regime (Figure 5B, bottom; Figure S8, middle).

Interestingly, a third circuit configuration likewise gave rise to excitatory and inhibitory neurons with matched classification accuracy near the biologically plausible regime (Figure 5C, bottom; Figure S8, right). In this configuration, inhibitory neurons were non-selective with respect to the excitatory pools, but were connected to the more selective excitatory neurons, i.e. those with a high signal-to-noise ratio (Figure 5C, top).

Our modeling results raise two questions. First, how can the inhibitory neurons have better decoding accuracy than the excitatory ones (Figure 5B,C, bottom; for part of the plot, red is above blue)? After all, in our model all information about the choice flows through the excitatory neurons. Second, why is the relative strength of the excitatory to inhibitory versus inhibitory to excitatory connections an important parameter (Figure 5, bottom; x-axis)? The answers are related. Increasing the strength of the excitatory to inhibitory connections increases the signal in the inhibitory neurons, and therefore effectively decreases the noise added to the inhibitory population (see Methods for details). This decrease in noise leads to improved decoding accuracy of both the excitatory and inhibitory populations, because the two populations are connected. However, the decrease in the noise added to the inhibitory neurons has a bigger effect on the inhibitory than the excitatory population; that’s because noise directly affects the inhibitory neurons, but only indirectly, through the inhibitory to excitatory connections, affects the excitatory neurons. Thus, in all panels of Figure 5, the classification accuracy increases faster for inhibitory neurons than excitatory ones as the excitatory to inhibitory connection strength increases.

Overall, our modeling work rules out decision circuits with non-selective inhibition (Figure 5A), and instead demonstrates that excitatory and inhibitory neurons in decision circuits must be selectively connected, either based on the signal preference (Figure 5B) or the informativeness (Figure 5C) of excitatory neurons.

### Correlations are stronger between similarly tuned neurons

We have demonstrated that inhibitory neurons are choice-selective (Figures 2,3). If choice selectivity in inhibitory neurons emerges because of functionally biased input from excitatory neurons, one prediction is that correlations will be stronger between excitatory and inhibitory neurons with the same choice selectivity compared to those with the opposite choice selectivity (Cossell et al., 2015; Francis et al., 2018). To test this hypothesis, we compared pairwise noise correlations in the activity of neurons with the same vs. opposite choice selectivity (Methods). Indeed, neurons with the same choice selectivity had stronger correlations (Figure 6A). This was evident in pairs consisting of one excitatory, one inhibitory, only excitatory, or only inhibitory neurons (Figure 6A, left to right), in keeping with previous observations in mouse V1 during passive viewing (Hofer et al., 2011; Ko et al., 2011; Cossell et al., 2015; Znamenskiy et al., 2018), as well as the prefrontal cortex in behaving monkeys (Constantinidis and Goldman-Rakic, 2002).

**Figure 6.**
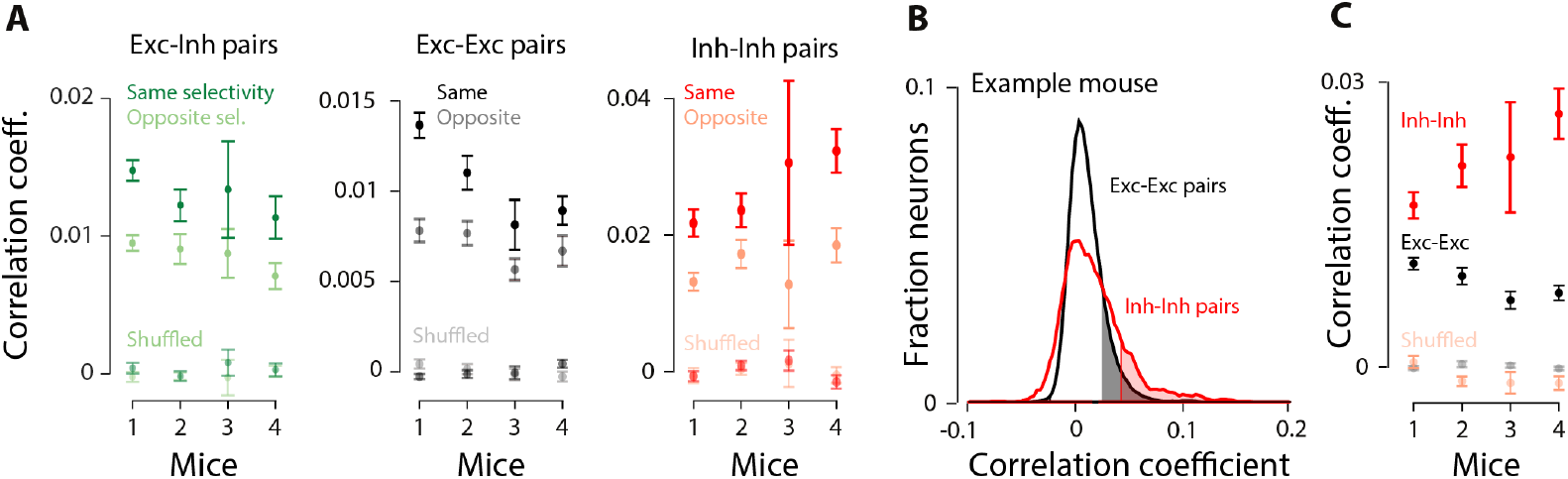
Pairwise noise correlations are stronger between neurons with the same choice selectivity. **A, Left:** Noise correlations (Pearson’s coefficient) for pairs of excitatory-inhibitory neurons with the same choice selectivity (dark green) or opposite choice selectivity (light green, i.e. one neuron prefers ipsilateral, and the other neuron prefers contralateral choice). **Middle, Right:** same as in the left panel, but for excitatory-excitatory, and inhibitory-inhibitory pairs, respectively. “Shuffled” denotes quantities were computed using shuffled trial labels. Mean+/-standard error across days; 0-97 ms before the choice. Same vs. opposite is significant in all cases, except for mouse 3 in EE and II pairs (t-test, p<0.05). **B**, Example mouse: distribution of noise correlations (Pearson’s correlation coefficients, 0-97 ms before the choice) for excitatory neurons (blue; n=11867) and inhibitory neurons (red; n=1583). Shaded areas indicate significant quantities compared to a shuffled control: trial orders were shuffled for each neuron to remove noise correlations. **C**, Summary of noise correlation coefficients for each mouse, indicating higher correlations among inhibitory neurons; mean+/-standard error across days.

The higher noise correlations among similarly tuned excitatory-inhibitory neuron pairs is also consistent with the observation that in V1, excitatory and inhibitory neurons that belong to the same subnetwork are reciprocally connected (Yoshimura and Callaway, 2005). An alternative explanation, that the neurons with similar tuning share common inputs, is also possible. However, these shared inputs are likely not sensory inputs because we observed the same correlation effects in the pre-trial period in which there is no stimulus (Figure S9A).

We next compared the strength of pairwise noise correlations within our excitatory and inhibitory populations. Inhibitory pairs had significantly higher noise correlations compared to excitatory pairs (Figure 6B,C: noise correlations; Figure S9C: spontaneous correlations). Importantly, we obtained the same results even when we restricted the analysis to those inhibitory and excitatory neurons that had the same inferred spiking activity (Figure S9D,E). This was done because the higher spiking activity of inhibitory neurons (Figure 1G-I) could potentially muddle the comparison of pairwise noise correlations between excitatory and inhibitory neurons. Finally, similar to previous reports (Hofer et al., 2011; Khan et al., 2018), we found intermediate correlations for pairs consisting of one inhibitory neuron and one excitatory neuron (Figure S9B,C). These findings align with previous studies in sensory areas that have demonstrated stronger correlations among inhibitory neurons (Hofer et al., 2011; Khan et al., 2018). These correlations are likely driven at least in part by local connections, as evidenced by the dense connectivity of interneurons with each other (Galarreta and Hestrin, 1999; Packer and Yuste, 2011; Kwan and Dan, 2012). The difference we observed between excitatory and inhibitory neurons argues that this feature of early sensory circuits is shared by decision-making areas. Further, this clear difference between excitatory and inhibitory neurons, like the difference in inferred spiking (Figure 1G-I) and outcome selectivity (Figure S6C), confirms that we successfully measured two distinct populations. Overall our noise correlation analysis suggests that selective connectivity between excitatory and inhibitory neurons exist and depends on their functional properties.

### Noise correlations limit decoding accuracy

Our results thus far demonstrate that neural activities in both excitatory and inhibitory populations reflect an animal’s impending choice (Figure 3B,C), and that there are significant noise correlations among neurons in PPC (Figure 6). However, the analyses so far do not demonstrate how this noise affects the ability to decode neural activity overall, or for excitatory and inhibitory populations separately. Examining the effect of noise is essential because noise correlations can limit or enhance the ability to decode population activity depending on their structure (Panzeri et al., 1999; Averbeck et al., 2006). Fortunately, our dataset includes simultaneous activity from hundreds of neurons and is therefore especially well-suited to assess noise correlations: correlations can have a large effect at the population level even when their effect at the level of neuron pairs is small (Averbeck et al., 2006; Moreno-Bote et al., 2014).

To examine how noise correlations affected classification accuracy for choice, we sorted neurons based on their individual choice selectivity, added them one by one to the population (from highest to lowest choice selectivity defined as |AUC-0.5|), and measured classification accuracy as a function of population size. Classification accuracy improved initially as more neurons were included in the decoder, but quickly saturated (Figure 7A black; 0-97 ms before the choice).

**Figure 7.**
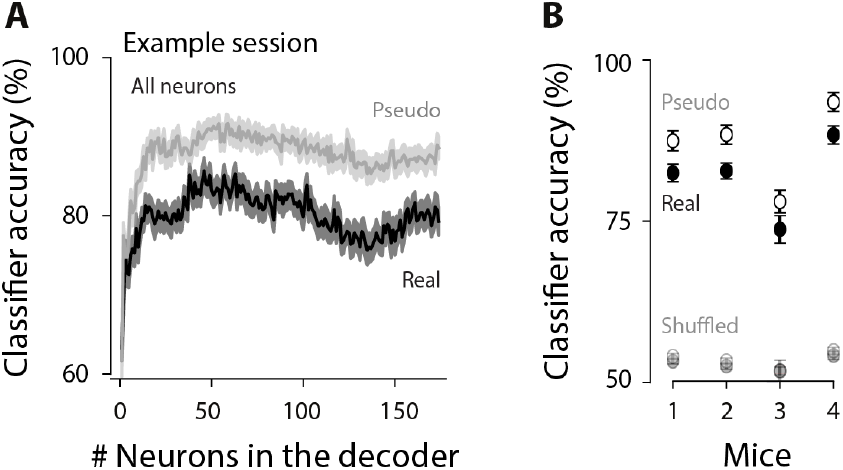
Noise correlations reduce classification accuracy. **A**, Classification accuracy for an example session (at time window 0-97 ms before the choice) on neural ensembles of increasingly larger size, with the most choice-selective neurons added first. Mean+/-standard error across 50 crossvalidated samples. Gray: classification accuracy for pseudopopulations, in which noise correlations were removed by shuffling. Black: real populations. Both cell types were included (“All neurons”). **B**, Summary for each mouse; points show mean+/-standard error across days. Values were computed for the largest neuronal ensemble (the max value on the horizontal axis in D).

To understand why classification accuracy saturates, we tested the effect of noise correlations on classification accuracy. Specifically, we created “pseudo populations”, in which each neuron in the population was taken from a different trial (Figure 7A gray). This removed noise correlations because those are shared across neurons within a single trial. Higher classification accuracy in pseudo populations compared to real populations indicates the presence of noise that overlaps with signal, limiting information (Panzeri et al., 1999; Averbeck et al., 2006; Averbeck and Lee, 2006; Moreno-Bote et al., 2014). This is what we observed (Figure 7A, gray trace above black trace). Across all mice, removing noise correlations resulted in a consistent increase in classification accuracy for the full population (Figure 7B; filled vs. open circles). This establishes that noise correlations limit population decoding in PPC.

### Selectivity increases in parallel in inhibitory and excitatory populations during learning

Our observations thus far argue that excitatory and inhibitory neurons form selective subnetworks. To assess whether the emergence of these subnetworks is experience-dependent, and if it varies between inhibitory and excitatory populations, we measured neural activity as animals transitioned from novice to expert decision-makers (3 mice; 35-48 sessions; Figure S10). We trained a linear classifier for each training session, and for each moment in the trial. This allowed us to compare the dynamics of the choice signal in excitatory and inhibitory populations over the course of learning.

Classification accuracy of the choice decoder increased consistently as animals became experts in decision-making (Figure 8A, left; Figure 8D, black), leading to a strong correlation between the classifier performance and the animal’s performance across training days (Figure 8B, left). The population representation of the choice signal also became more prompt: the choice signal appeared progressively earlier in the trial as the animals became experts. Initially, classification accuracy was high only after the choice (Figure 8A, black arrow). As the animals gained experience, high classification accuracy occurred progressively earlier in the trial, eventually long before the choice (Figure 8A, gray arrow). This resulted in a negative correlation between the animal’s performance and the onset of super-threshold decoding accuracy relative to the choice (Figure 8C, left; Figure 8E, black).

**Figure 8.**
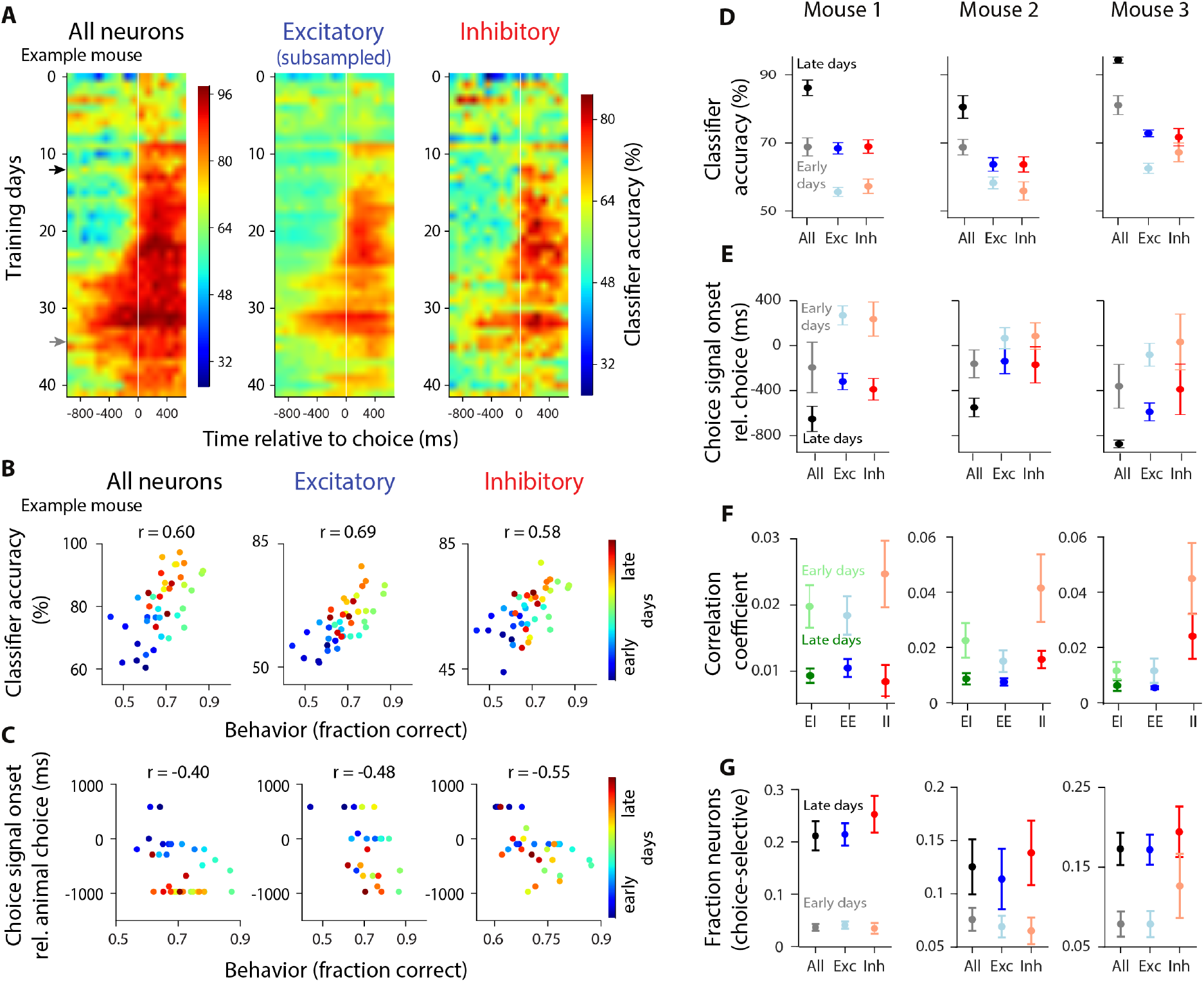
Learning leads to increased magnitude of the choice signal in the population, increased fraction of choice-selective neurons, and reduced noise correlations, in both excitatory and inhibitory populations. **A**, Decoder accuracy is shown for each training session, for all neurons (left), subsampled excitatory (middle), and inhibitory neurons (right). White vertical line: choice onset. Each row: average across cross-validation samples; example mouse. Colorbar of the inhibitory plot applies to the excitatory plot too. **B**, Scatter plot of classifier accuracy at 0-97 ms before the choice vs. behavioral performance (fraction correct on easy trials), including all training days. r is Pearson correlation coefficient (p<0.001 in all plots); same example mouse as in (A). Correlations for behavior vs. classification accuracy for all neurons, excitatory and inhibitory: 0.55, 0.35, 0.32 in mouse 2; 0.57, 0.63, 0.32 in mouse 3. Correlations for behavior vs. choice-signal onset for all neurons, excitatory and inhibitory: −0.60, −0.34, −0.38, in mouse 2; −0.60, −0.27, −0.28 in mouse 3. All values: p<0.05 **C**, Same as (B), except showing the onset of choice signal, i.e. the first moment in the trial that classifier accuracy was above chance (ms, relative to choice onset) vs. behavioral performance. **D**, Summary of each mouse, showing classification accuracy averaged across early (dim colors) vs. late (dark colors) training days. **E**, Same as (D), but showing choice signal onset (ms). **F**, Same as (D), but showing pairwise noise correlation coefficients. **G**, Fraction of choice-selective neurons increases as a result of training; average across early (dim colors) and late (dark colors) training days; time points 0-97 ms before the choice. Early days were the first few training days in which the animal’s performance was lower than the 20th percentile of animal’s performance across all days. Late days included the last training days in which the animal’s behavioral performance was above the 80th percentile of performance across all days.

Importantly, the dynamics of the choice decoder changed in parallel in both excitatory and inhibitory populations as a result of training: the choice signal emerged at the same time in both populations, and its magnitude and timing was matched for the two cell types throughout learning (Figure 8A-C, middle, right; Figure 8D-E, blue, red). This change was not due to the presence of more correct trials in later sessions: an improvement in classification accuracy was clear even when the number of correct trials was matched for each session (Figure S12C). These findings indicate that learning induces the simultaneous emergence of choice-specific subpopulations in excitatory and inhibitory cells in PPC.

Notably, the animal’s licking or running behavior could not explain the learning-induced changes in the magnitude of classification accuracy (Figure S11). The center-spout licks that preceded the left vs. right choices were overall similar during the course of learning (Figure S11A), and did not differ in early vs. late training days (Figure S11B). The similarity in lick movements for early vs. late sessions stands in contrast to the changes in the classification accuracy for early vs. late sessions (Figure 8). We also assessed animals’ running behavior during the course of learning (Figure S11C,D). In some sessions, the running distance differed preceding left vs. right choices (Figure S11C). Nonetheless, when we restricted our analysis to days in which the running distance was indistinguishable for the two choices (0-97 ms before the choice, t-test, P>0.05), we were still able to accurately classify the animal’s choice using neural activity (Figure S11D). These observations provide reassurance that the population activity does not entirely reflect preparation of licking and running movements, and argue instead that the population activity reflects the animal’s stimulus-informed choice.

Finally, we studied how cofluctuations changed over the course of training. Pairwise correlations in neural activity were overall higher in early training days, when mice were novices, compared to late training days, as they approached expert behavior (Figure 8F, unsaturated colors above saturated colors). This effect was observed for all combinations of neural pairs (Figure 8F, green: excitatory-inhibitory; blue: excitatory-excitatory; red: inhibitory-inhibitory). These findings are in agreement with previous reports suggesting that learning results in reduced noise correlations (Gu et al., 2011; Jeanne et al., 2013; Khan et al., 2018; Ni et al., 2018), enhancing information that is encoded in neural populations. To test if the learning-induced increase in classification accuracy (Figure 8A,B,D) was entirely a consequence of the reduction in noise correlations (Figure 8F), we studied how classification accuracy of pseudo populations, which lack noise correlations, changed with training. Interestingly, we still observed a significant increase in the classification accuracy of pseudo populations as a result of training (Figure S12A,B). Therefore, the reduction in noise correlations cannot alone account for the improved classification accuracy that occurs during learning. Instead, it suggests that choice selectivity of individual neurons also changes with learning. Indeed, the fraction of choice-selective neurons increased threefold, in both excitatory and inhibitory cell types, as a result of training (Figure 8G), contributing to the improved classification accuracy at the ensemble level.

## Discussion

Despite a wealth of studies assessing the selectivity of inhibitory neurons in response to sensory features, little is known about the selectivity of inhibitory neurons in decision-making. This represents a critical gap in our knowledge, and has left untested key features of decision-making models relying on inhibitory neurons. To close this gap, we simultaneously measured excitatory and inhibitory populations during perceptual decisions about multisensory stimuli.

We demonstrated that excitatory and inhibitory neurons predict the animal’s impending choice with equal fidelity (Figure 2,3). This result, along with our modeling (Figure 5), constrains circuit models of decision-making, ruling out models in which inhibitory neurons receive completely nonspecific input from excitatory populations (Figure 5A). Instead, our findings suggest that specific functional subnetworks exist within inhibitory populations, just like excitatory populations (Figure 5B). This implies targeted connectivity between excitatory and inhibitory neurons (Yoshimura and Callaway, 2005; Znamenskiy et al., 2018), and supports circuit architectures with functionally specific subnetworks within excitatory and inhibitory populations that are reciprocally connected.

The advantage of signal-selective architecture is that it offers improved stability (Znamenskiy et al., 2018) and robustness to perturbations (Lim and Goldman, 2013). In a recent study (Lim and Goldman, 2013), candidate circuit architectures were subjected to small perturbations that are likely to occur in real brains, such as changes in the network’s intrinsic gain, loss of excitatory/inhibitory neurons, changes in the strengths of excitatory/inhibitory synaptic transmission, and global shifts in background input. The intuition is that negative feedback from a specific pool can oppose drifts resulting from these changes, allowing the network to remain stable. In the absence of such correction, the network can easily become unstable even after a fairly minor perturbation (e.g., a 1% increase in intrinsic gain,–Lim & Goldman, Fig 6, j-l). Another recent study (Znamenskiy et al., 2018) suggested that targeted connectivity between excitatory and inhibitory neurons allows for the existence of highly selective excitatory subnetworks, while keeping the network stable. In circuits with non-selective inhibition, by contrast, excitatory subnetworks had to be weakly selective in order to keep the network stable (Znamenskiy et al., 2018). The permissiveness to highly selective excitatory subnetworks in circuits with selective inhibition may be advantageous for situations that require precise encoding of sensory stimuli for discrimination.

The stability and robustness of specific inhibition models discussed above are appealing for decision-making. However, those studies (Lim and Goldman, 2013) did not aim to describe behavior and electrophysiological responses during decision-making in nearly the detail of previous studies that leveraged traditional, non-selective inhibition (Wang, 2002; Bogacz et al., 2006; Wong and Wang, 2006). Those traditional models accurately captured numerous features of evidence integration, leaving it to the experimentalists to assess their implementation at the circuit level. However, the nonspecific inhibition in traditional implementations does not agree with the experimental data reported here and thus must be revisited. We propose an alternative circuit model that relies on specific connectivity between excitatory and inhibitory neurons, determined by signal preference (Figure 5B). Evaluating the performance of this revised model in predicting decision-making behavior and neural activity can help further constrain its implementation. Additionally, it will generate new predictions that can subsequently be evaluated at the circuit level. Examples include predictions about the strength of connections between neural populations, their dependence on signal and noise, tuning of the network to distinct task components, and network modifications during learning.

The equal selectivity for choice that we observed in excitatory and inhibitory populations is surprising: the broad stimulus tuning curves observed in most V1 inhibitory neurons (Sohya et al., 2007; Niell and Stryker, 2008; Kerlin et al., 2010; Bock et al., 2011; Hofer et al., 2011; Znamenskiy et al., 2018) (but see Runyan et al., 2010) and the dense connectivity for inhibitory neurons (Hofer et al., 2011; Packer and Yuste, 2011; Znamenskiy et al., 2018) are often taken as evidence that inhibitory neurons are not strongly modulated by task parameters. Two differences between our study and previous ones may explain why we saw equal selectivity in excitatory and inhibitory populations.

First, we measured neural activity in PPC where the proportion of interneuron subtypes differ from V1; in particular, early sensory areas are more enriched in PV interneurons relative to SOM and VIP neurons, whereas the opposite is true in association areas (Kim et al., 2017; Wang and Yang, 2018). Moreover, interneuron subtypes vary in their specificity of connections (Pfeffer et al., 2013); for instance, PV interneurons are suggested to have broader tuning than SOM and VIP cells (Wang et al., 2004; Ma et al., 2010). Therefore, the strong selectivity that we found in all GABAergic interneurons in PPC may not contradict the broad selectivity observed in studies largely performed on PV interneurons in V1. Future studies that measure the selectivity of distinct interneuron populations during decision-making in V1 vs. PPC will be helpful. Here, we measured all GABAergic interneurons instead of individual interneuron subtypes; this was because of the technical challenges in reliably identifying more than two cell types in a single animal, and because of the importance of simultaneously measuring the activity of excitatory and inhibitory neurons within the same subject. Had we lacked within-animal measurements, our ability to compare excitatory vs. inhibitory neurons would have been compromised by animal-to-animal variability (e.g. note the matched selectivity of excitatory and inhibitory neurons within each subject in Figure 3C despite the overall variability in selectivity across subjects).

Second, analyzing neural activity in the context of decision-making naturally led us to make different comparisons than those carried out in previous work. For example, we measured selectivity for a binary choice, while sensory tuning curves are measured in response to continuously varying stimuli (e.g., orientation). Further, we measured activity in response to an abstract stimulus, the meaning of which was learned gradually by the animal. This may recruit circuits that differ from those supporting sensory processing in passively viewing mice. Finally, we used stochastically fluctuating multisensory stimuli, which have not been evaluated in mouse V1. Future studies that examine the tuning of V1 neurons to the sensory stimulus used here will determine if V1 inhibitory neurons will be as sharply tuned as excitatory neurons to the stimulus. This is a possibility: the tuning strength of interneurons can vary substantially for different stimulus features. For instance, PV neurons in V1 have particularly poor tuning to the orientation of visual stimuli, while their temporal-frequency tuning is considerably stronger (Znamenskiy et al., 2018).

Our long-term monitoring of neural activity within the same subjects provides a critical new insight into decision-making circuitry by demonstrating how acquiring expertise modulates the activity of excitatory and inhibitory neurons in PPC. We observed that learning induced an increase in the number of choice-selective neurons and a decrease in noise correlations, indicating plasticity and reorganization of connections. As a result, population responses preceding the two choices became progressively more distinct with training. Importantly, these changes occurred in parallel in both excitatory and inhibitory cells. Our findings are partially in agreement with those in V1, in which learning improves tuning to sensory stimuli in excitatory (Schoups et al., 2001; Poort et al., 2015; Khan et al., 2018) and some inhibitory subtypes (Khan et al., 2018). However, in V1 excitatory neurons have stronger tuning to sensory stimuli early in training (Khan et al., 2018); in contrast, the magnitude of choice selectivity in PPC was the same for both cell types throughout training in our study (Figure 8). Primate studies have likewise observed that perceptual learning changes the selectivity of neurons (Freedman and Assad, 2006; Law and Gold, 2008; Viswanathan and Nieder, 2015) and reduces noise correlations (Gu et al., 2011; Ni et al., 2018).

Finally, we demonstrated that the learning-induced changes in PPC selectivity were closely associated with the changes in animal performance, in keeping with primate studies of decision-making (Law and Gold, 2008). This, together with our finding that changes in population activity do not purely reflect movements (Figure S11), further corroborates the suggested role for PPC in mapping sensation to action (Law and Gold, 2008; Raposo et al., 2014; Pho et al., 2018). Future experiments using causal manipulations will reveal whether the increased choice selectivity we observed in PPC originates there or is inherited from elsewhere in the brain.

By measuring cell-type-specific activity in parietal cortex during decision-making, we have provided evidence that excitatory and inhibitory populations are equally choice-selective, and that these ensembles emerge in parallel, as mice become skilled decision-makers. These results argue against models with non-specific connectivity between excitatory and inhibitory neurons, at least in decision circuits. In future modeling efforts, these features can be incorporated into decision-making models, and their impact on key model outputs, e.g. reaction time distributions and firing rates, can be evaluated. Such studies will shed light on how microcircuits of inhibitory and excitatory neurons may vary across areas in their selectivity and specificity of connections, and will reveal the circuit architectures that allow for equally selective inhibitory and excitatory neurons.

## Methods

### Imaging and behavioral dataset

Our simultaneous imaging and decision-making dataset includes 135 sessions from 4 mice (45, 48, 7, and 35 sessions per mouse). Median number of trials per session is 213, 253, 264, and 222, for each mouse. On average, 480 neurons were imaged per session, out of which ~40 neurons were inhibitory and ~330 were excitatory. Approximately 100 neurons per session were not classified as either excitatory or inhibitory since they did not meet our strict cell-type classification criteria (see below). In 3 of the mice, the same group of neurons was imaged throughout learning (35-48 training days).

### Mice and surgical procedure

Gad2-IRES-CRE (Taniguchi et al., 2011) mice were crossed with Rosa-CAG-LSL-tdTomato-WPRE (aka Ai14; Madisen et al., 2010) to create mice in which all GABAergic inhibitory neurons were labeled. Adult mice (~2-month old) were used in the experiments. Meloxicam (analgesic), dexamethasone (anti-inflammatory) and Baytril (enrofloxacin; anti-biotic) were injected 30min before surgery. Using a biopsy punch, a circular craniotomy (diameter: 3mm) was made over the left PPC (stereotaxic coordinates: 2 mm posterior, 1.7 mm lateral of bregma (Harvey et al., 2012) under isoflurane (~5%) anesthesia. Pipettes (10-20 um in diameter, cut at an angle to provide a beveled tip) were front-filled with AAV9-Synapsin-GCaMP6f (U Penn, Vector Core Facility) diluted 2X in PBS (Phosphae-buffered saline). The pipette was slowly advanced into the brain (Narishige MO-8 hydraulic micro-manipulator) to make ~3 injections of 50nL, slowly over an interval of ~5-10 min, by applying air pressure using a syringe. Injections were made near the center of craniotomy at a depth of 250-350 μm below the dura. A glass plug consisting of a 5mm coverslip attached to a 3mm coverslip (using IR-curable optical bond, Norland) was used to cover the craniotomy window. Vetbond, followed by metabond, was used to seal the window. All surgical and behavioral procedures conformed to the guidelines established by the National Institutes of Health and were approved by the Institutional Animal Care and Use Committee of Cold Spring Harbor Laboratory.

### Imaging

We used a 2-photon Moveable Objective Microscope with resonant scanning at approximately 30 frames per second (Sutter Instruments, San Francisco, CA). A 16X, 0.8 NA Nikon objective lens was used to focus light on fields of view of size 512−512 pixels (~575 μm x ~575 μm). A Ti:sapphire laser (Coherent) delivered excitation light at 930nm (average power: 20-70 mW). Red (ET670/50m) and green (ET 525/50m) filters (Chroma Technologies) were used to collect red and green emission light. The microscope was controlled by Mscan (Sutter). In mice in which chronic imaging was performed during learning, the same plane was identified on consecutive days using both coarse alignment, based on superficial blood vessels, as well as fine alignment, using reference images of the red channel (tdTomato expression channel) at multiple magnification levels. For each trial, imaging was started 500ms before the trial-initiation tone, and continued 500ms after reward or time-out. We aimed to image in the center of the window for all mice, but in one animal (Mouse 4), some tissue regrowth obscured the signal in this region and so imaging was performed slightly further back.

### Decision-making behavior

Mice were gradually water restricted over the course of a week, and were weighed daily. Mice harvested at least 1 mL of water per behavioral/imaging session, and completed 100-500 trials per session. After approximately one week of habituation to the behavioral setup, 15-30 training days were required to achieve 75% correct choice. Animal training took place in a sound isolation chamber. The stimulus in all trials was multisensory, consisting of a series of simultaneous auditory clicks and visual flashes, occurring with Poisson statistics (Brunton et al., 2013; Odoemene et al., 2017). Multisensory stimuli were selected because they increased the learning rate of the mice, a critical consideration since GCaMP6f expression can be unreliable over a long period of time. Stimulus duration was 1000 ms. Each pulse was 5 ms; minimum interval between pulses was 32 ms, and maximum interval was 250 ms. The pulse rate ranged from 5 to 27 Hz. The category boundary for marking high-rate and low-rate stimuli was 16 Hz, at which animals were rewarded randomly on either side. The highest stimulus rates used here are known to elicit reliable, steady state flicker responses in retinal ERG in mice (Krishna et al., 2002; Tanimoto et al., 2015).

Mice were on top of a cylindrical wheel and a rotary encoder was used to measure their running speed. Trials started with a 50 ms initiation tone (Figure S1A). Mice had 5 sec to initiate a trial by licking the center waterspout (Marbach and Zador, 2017), after which the multisensory stimulus was played for 1 second. If mice again licked the center waterspout, they received 0.5 μL water on the center spout, and a 50ms go cue was immediately played. Animals had to report a choice by licking to the left or right waterspout within 2 sec. Mice were required to confirm their choice by licking the same waterspout one more time within 300 ms after the initial lick (Marbach and Zador, 2017). The “confirmation lick” helped dissociate the choice time (i.e. the time of first lick to the side waterspout), from the reward time (i.e. the time of second lick to the side waterspout); it also helped with reducing impulsive choices. If the choice was correct, mice received 2-4 μL water on the corresponding waterspout. An incorrect choice was punished with a 2 sec time-out. The experimenter-imposed inter-trial intervals (ITI) were drawn from a truncated exponential distribution, with minimum, maximum, and lambda equal to 1 sec, 5 sec, and 0.3 sec, respectively. However, the actual ITIs could be much longer depending on when the animal initiates the next trial. Bcontrol (Raposo et al., 2014) with a Matlab interface was used to deliver trial events (stimulus, reward, etc) and collect data.

### Logistic regression model of behavior

A modified version of the logistic regression model in (Busse et al., 2011) was used to assess the extent to which the animal’s choice depends on the strength of sensory evidence, i.e. how far the stimulus rate is from the category boundary at 16Hz, the previous choice outcome (success or failure) and ITI, i.e. the time interval between the previous choice and the current stimulus onset (Figure S1B).

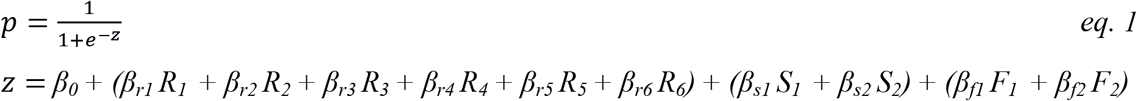

where *p* is the probability of choosing the left choice, and *z* is the decision variable. *R*, *S* and *F* are vectors of indicator variables; each element corresponds to 1 trial. Stimulus strength (*R*) was divided into 6 bins (*R_1_* to *R_6_*). Previous success (*S*) was divided into 2 bins (*S_1_* to *S_2_*): success after a long ITI (> 7sec) and success after a short ITI (< 7sec). Previous failure (*F*) was divided into 2 bins (*F_1_* to *F_2_*): failure after a long and short ITI. For instance, if a trial had stimulus strength 3 Hz, and was preceded by a success choice with ITI 5 sec, then R_2_ and S_1_ would be set to 1 and all other R, S and F parameters to 0 (Figure S1B).

For each session the scalar coefficients *β_0_, β_r1_* to *β_r6_, β_s1_, β_s2_, β_f1_*, and *β_f2_* were fitted using Matlab glmfit.m. Figure S1B left shows *β_r1_* to *β_r6_*. Figure S1B middle shows *β_s1_* and *β_s2_*, and Figure S1B right shows *β_f1_* and *β_f2_*.

### ROI (region of interest) extraction and deconvolution

The recorded movies from all trials were concatenated and corrected for motion artifacts by cross-correlation using DFT registration (Guizar-Sicairos et al., 2008). Subsequently, active ROIs (sources) were extracted using the CNMF algorithm (Pnevmatikakis et al., 2016) as implemented in the CaImAn package (Giovannucci et al., 2019) in MATLAB. The traces of the identified neurons were ΔF/F normalized and then deconvolved by adapting the FOOPSI deconvolution algorithm (Vogelstein et al., 2010; Pnevmatikakis et al., 2016) to a multi-trial setup. This was necessary because simply concatenating individual trials would lead to discontinuities in the traces, which could distort estimates of the time constants. Each value of Foopsi deconvolution represents spiking activity at each frame for a given neuron. We have referred to the deconvolved values as “inferred spiking activity” throughout the paper. The deconvolved values do not represent absolute firing rates, so they cannot be compared across neurons. However, for a particular neuron, higher inferred spiking activity means higher firing rate. We elected to base our analyses on inferred spikes rather than fluorescence activity because peak amplitudes and time constants of the fluorescence responses vary across neurons, affecting subsequent analyses (Machado et al., 2015; Helmchen and Tank, 2019).

The adaptation of the FOOPSI for multi-trial setup involved the following steps. For each component, the activity trace over all the trials was used to determine the time constants of the calcium indicator dynamics as in (Pnevmatikakis et al., 2016). Then the neural activity during each trial was deconvolved separately using the estimated time constant and a zero baseline (since the traces were ΔF/F normalized). A difference of exponentials was used to simulate the rise and decay of the indicator.

### Neuropil Contamination removal

The CNMF algorithm demixes the activity of overlapping neurons. It takes into account background neuropil activity by modeling it as a low rank spatiotemporal matrix (Pnevmatikakis et al., 2016). In this study a rank two matrix was used to capture the neuropil activity. To evaluate its efficacy we compared the traces obtained from CNMF to the traces from a “manual” method similar to (Chen et al., 2013) (Figure S13): the set of spatial footprints (shapes) extracted from CNMF were binarized by thresholding each component at the 0.2x its maximum value level. The binary masks were then used to average the raw data and obtain an activity trace that also included neuropil effects. To estimate the background signal, an annulus around the binary mask was constructed with minimum distance 3 pixels from the binary mask and width 7 pixels (Figure S13A). The average of the raw data over the annulus defined the background trace, which was then subtracted from the activity trace. The resulting trace was then compared with the CNMF estimated temporal trace for this activity. The comparison showed a very high degree of similarity between the two traces (Figure S13; example component; r=0.96), with the differences between the components being attributed to noise and not neuropil related events. Note that this “manual” approach is only applicable in the case when the annulus does not overlap with any other detected sources. These results demonstrate the ability of the CNMF framework to properly capture neuropil contamination and remove it from the detected calcium traces.

### ROI inclusion criteria

We excluded poor-quality ROIs identified by the CNMF algorithm based on a combination of criteria: 1) size of the spatial component, 2) decay time constant, 3) correlation of the spatial component with the raw ROI image built by averaging spiking frames, 4) correlation of the temporal component with the raw activity trace, and 5) the probability of fluorescence traces maintaining values above an estimated signal-to-noise level for the expected duration of a calcium transient(Giovannucci et al., 2018) (GCaMP6f, frame rate: 30Hz). A final manual inspection was performed on the selected ROIs to validate their shape and trace quality.

### Identification of inhibitory neurons

We used a two-step method to identify inhibitory neurons. First, we corrected for bleed-through from green to red channel by considering the following regression model,

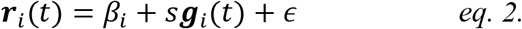

where, ***r**_i_*(*t*) and ***g**_i_*(*t*) are vectors, indicating pixel intensity in red and green channel, respectively, with each component of the vector corresponding to one pixel in the ROI. i labels ROI (presumably each ROI is a neuron). *β_i_* is the offset, and s is the parameter that tells us how much of the green channel bleeds through to the red one. 1_*p*_ ∈ ℝ^*p*^ is a vector whose components are all 1.

It is the parameter *s* that we are interested in. To find *s*, we define a cost function, *C*,

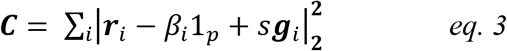

and minimize it with respect to s and all the *β_i_*. The value of s at the minimum reflects the fraction of bleed-through from the green to the red channel. That value, denoted s*, is then used to compute the bleedthrough-corrected image of the red-channel, denoted *I* via the expression

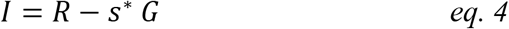

where *R* and *G* are the time-averaged images of the red and green channels, respectively.

Once the bleedthrough-corrected image, *I*, was computed, we used it to identify inhibitory neurons using two measures:

1. A measure of local contrast, by computing, on the red channel, the average pixel intensity inside each ROI mask relative to its immediate surrounding mask (width=3 pixels). Given the distribution of contrast levels, we used two threshold levels, *T_E_* and *T_I_*, defined, respectively, as the 80^th^ and 90^th^ percentiles of the local contrast measures of all ROIs. ROIs whose contrast measure fell above *T_I_* were identified as inhibitory neurons. ROIs whose contrast measure fell below *T_E_* were identified as excitatory neurons, and ROIs with the contrast measure in between *T_E_* and *T_I_* were not classified as either group (“unsure” class).
2. In addition to a measure of local contrast, we computed for each ROI the correlation between the spatial component (ROI image on the green channel) and the corresponding area on the red channel. High correlation values indicate that the ROI on the green channel has a high signal on the red channel too; hence the ROI is an inhibitory neuron. We used this correlation measure to further refine the neuron classes computed from the local contrast measure (i.e. measure 1 above). ROIs that were identified as inhibitory based on their local contrast (measure 1) but had low red-green channel correlation (measure 2), were reset as “unsure” neurons. Similarly, ROIs that were classified as excitatory (based on their local contrast) but had high red-green channel correlation were reclassified as unsure. Unsure ROIs were included in the analysis of all-neuron populations (Figure 3A left); but were excluded from the analysis of excitatory only or inhibitory only populations (Figure 3A middle, right). Finally, we manually inspected the ROIs identified as inhibitory to confirm their validity. This method resulted in 11% inhibitory neurons, which is within the range of previous studies (10-20%: Rudy et al., 2011); (15%: Beaulieu, 1993); (16%: Gabbott et al., 1997); (<5%: de Lima and Voigt, 1997); (10-25%: de Lima et al., 2009).

### General analysis procedures

All analyses were performed on inferred spiking activity. Traces were down-sampled, so each bin was the non-overlapping moving average of 3 frames (97.1 ms). Inferred spiking activity for each neuron was normalized so the max spiking activity for each neuron equaled 1. The trace of each trial was aligned to the time of the choice (i.e. the time of the 1^st^ lick to either of the side waterspouts after the go tone). Two-tailed t-test was performed for testing statistical significance. Summary figures including all mice were performed on the time bin preceding the choice, i.e. 0-97 ms before choice onset. All reported correlations are Pearson’s coefficients. Analyses were performed in Python and Matlab.

### ROC analysis

The area under the ROC curve (AUC) was used to measure the choice preference of single neurons. Choice selectivity was defined as the absolute deviation of AUC from chance level (0.5). To identify significantly choice-selective neurons, for each neuron we performed ROC on shuffled trial labels (i.e. left and right choices were randomly assigned to each trial). This procedure was repeated 50 times to create a distribution of shuffled AUC values for each neuron (Figure S3A, “shuffled”). A neuron’s choice selectivity was considered to be significant if the probability of the actual AUC (Figure S3A, “real”) being drawn from the shuffled AUC distribution was less than 0.05. Time points from 0–97 ms before the decision were used to compute the fraction of choice-selective neurons (Figure 2B; Figure 8G).

### Decoding population activity

A linear SVM (Python sklearn package) was trained on each bin of the population activity in each session (non-overlapping 97ms time bins). To break any dependencies on the sequence of trials, we shuffled the order of trials for the entire population. To avoid bias in favor of one choice over the other, we matched the number of left- and right-choice trials used for classifier training. L2 regularization was used to avoid over-fitting. 10-fold cross validation was performed by leaving out a random 10% subset of trials to test the classifier performance, and using the remaining trials for training the classifier. This procedure was repeated 50 times. A range of regularization values was tested, and the one that gave the smallest error on the validation dataset was chosen as the optimal regularization parameter. Classifier accuracy was computed as the percentage of testing trials in which the animal’s choice was accurately predicted by the classifier, and summarized as the average across the 50 repetitions of trial subsampling. A minimum of 10 correct trials per choice was required in order to run the SVM on a session. Inferred spiking activity of each neuron was z-scored before running the SVM.

When comparing classification accuracy for excitatory vs. inhibitory neurons, the excitatory population was randomly sub-sampled to match the population size of inhibitory neurons to enable a fair comparison (Figure 3, blue vs. red). To compare the distribution of weights in the all-neuron classifier (Figure 3 black), the weight vector for each session was normalized to unity length (Figure 3D-F).

When decoding the stimulus category (Figure S6B), we used stimulus-aligned trials, and avoided any contamination by the choice signal by sub-selecting equal number of left and right choice trials for each stimulus category. When decoding trial outcome (Figure S6C), we used outcome-aligned trials, and avoided contamination by the choice or stimulus signal by subselecting equal number of trials from left and right choice trials for each trial outcome.

### Stability

To test the stability of the population code, decoders were trained and tested at different time bins (Kimmel et al., 2016) (Figure 4). To avoid the potential effects of auto-correlation, we performed cross validation not only across time bins, but also across trials. In other words, even though the procedure was cross validated by testing the classifier at a time different from the training time, we added another level of cross-validation by testing on a subset of trials that were not used for training. This strict method allowed our measure of stability duration to be free of auto-correlation effects.

As an alternative measure of stability, the angle between pairs of classifiers that were trained at different moments in the trial was computed (Figure S9C). Small angles indicate alignment, hence stability, of the classifiers. Large angles indicate misalignment, i.e. instability of the classifiers.

### Noise correlations

To estimate noise correlations at the population level, the order of trials was shuffled for each neuron independently. Shuffling was done within the trials of each choice, hence retaining the choice signal, while de-correlating the population activity to remove noise correlations. Then we classified population activity in advance of left vs. right choice (at time bin 0–97 ms before the choice) using the de-correlated population activity. This procedure was performed on neural ensembles of increasingly larger size, with the most selective neurons (|AUC-0.5|) added first (Figure 7A). To summarize how noise correlations affected classification accuracy in the population (Figure 7B), we computed, for the largest neural ensemble (Figure 7A, max value on the horizontal axis), the change in classifier accuracy in the de-correlated data (“pseudo populations”) vs. the original data. This analysis was only performed for the entire population; the small number of inhibitory neurons in each session prevented a meaningful comparison of classification accuracy on real vs. pseudo populations.

To measure pairwise noise correlations, we subtracted the trial-averaged response to a particular choice from the response of single trials of that choice. This allowed removing the effect of choice on neural responses. The remaining variability in trial-by-trial responses can be attributed to noise correlations, measured as the Pearson correlation coefficient for neuron pairs. We also measured noise correlations using the spontaneous activity defined as the neural responses in 0-97 ms preceding the trial initiation tone (Figure S9A,C). We computed the pairwise correlation coefficient (Pearson) for a given neuron with each other neuron within an ensemble (e.g., excitatory neurons). The resulting coefficients were then averaged to generate a single correlation value for that neuron. This was repeated for all neurons within the ensemble (Figure 6).

To compute pairwise correlations on excitatory and inhibitory neurons with the same inferred spiking activity (Figure S9D,E), we computed the median inferred spiking activity across trials for individual excitatory and inhibitory neurons in a session. The medians were then divided into 50 bins. The firing-rate bin that included the maximum number of inhibitory neurons was identified (“max bin”); inhibitory and excitatory neurons whose firing rate was within this “max bin” were used for the analysis. The firing rates were matched for these neurons because their median firing rate was within the same small bin of firing rates. Pairwise correlations were then computed as above.

### Learning analysis

In 3 of the mice, the same field of view was imaged each session during learning. This was achieved in two ways. First, the vasculature allowed a coarse alignment of the imaging location from day to day. Second, the image from the red channel was used for a finer alignment. Overall, most neurons were stably present across sessions (Figure S10). This suggests that we likely measured activity from a very similar population each day. Importantly, however, our conclusions do not rely on this assumption: our measures and findings focus on learning-related changes in the PPC population overall, as opposed to tracking changes in single neurons. To assess how population activity changed over learning, we evaluated classification accuracy each day, training a new decoder for each session. This approach allowed us to compute the best decoding accuracy for each session.

“Early days” (Figure 8; Figures S11,S12) included the initial training days in which the animal’s performance, defined as the fraction of correct choices on easy trials, was lower than the 20th percentile of performance across all days. “Late days” (Figure 8; Figures S11,S12) included the last training days in which the animal’s behavioral performance was above the 80^th^ percentile of performance across all days.

To measure the timing of decision-related activity (Figure 8C,E), we identified all sessions in which classifier accuracy was significantly different than the shuffle (t-test, p<0.05) over a window of significance that was at least 500 ms long. We defined the “choice signal onset” (Figure 8C,E) as the trial time corresponding to the first moment of that window. Sessions in which the 500 ms window of significance was present are included in Figure 8C. The number of points (and hence the relationship between session number and color in Figure 8C) differs slightly across the three groups. This is because on some sessions, the window of significance was present in one group but not another. For example, in a session the population including all neurons might have a 500 ms window of significance, hence it will contribute a point to Figure 7C left, while the population with only inhibitory neurons might be only transiently significant for <500ms, hence it will be absent from Figure 8C right.

### Modeling decision circuits

We considered a linearized rate network of the form

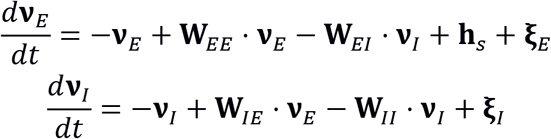

where *E* and *I* refer to the excitatory and inhibitory populations, respectively, **ν**_*E*_ and **ν**_*I*_ are vectors of firing rates (**ν**_*E*_ = *ν*_*E*1_, *ν*_*E*2_, …, and similarly for **ν**_*I*_), **W_*EE*_**, **W_*EI*_**, **W_*IE*_** and **W_*II*_** are the connectivity matrices (**W_*EI*_** indicates connection from inhibitory to excitatory neuron). **h**_*s*_ is the input, with s either 1 or 2 (corresponding to left and right licks), and **ξ** is trial to trial noise, taken to be zero mean and Gaussian, with covariance matrices

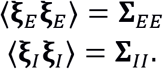

For the input we’ll assume that about half the elements of h_*s*_ are *h*_0_ for the rightward choice and −*h*_0_ for the leftward choice, and the rest are − *h*_0_ for the rightward choice and *h*_0_ for the leftward choice. We used *h*_0_ = 0.1 (see Table 1). The noise covariance is diagonal but non-identity, with diagonal elements distributed as

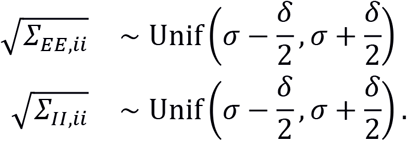

**Table 1.**
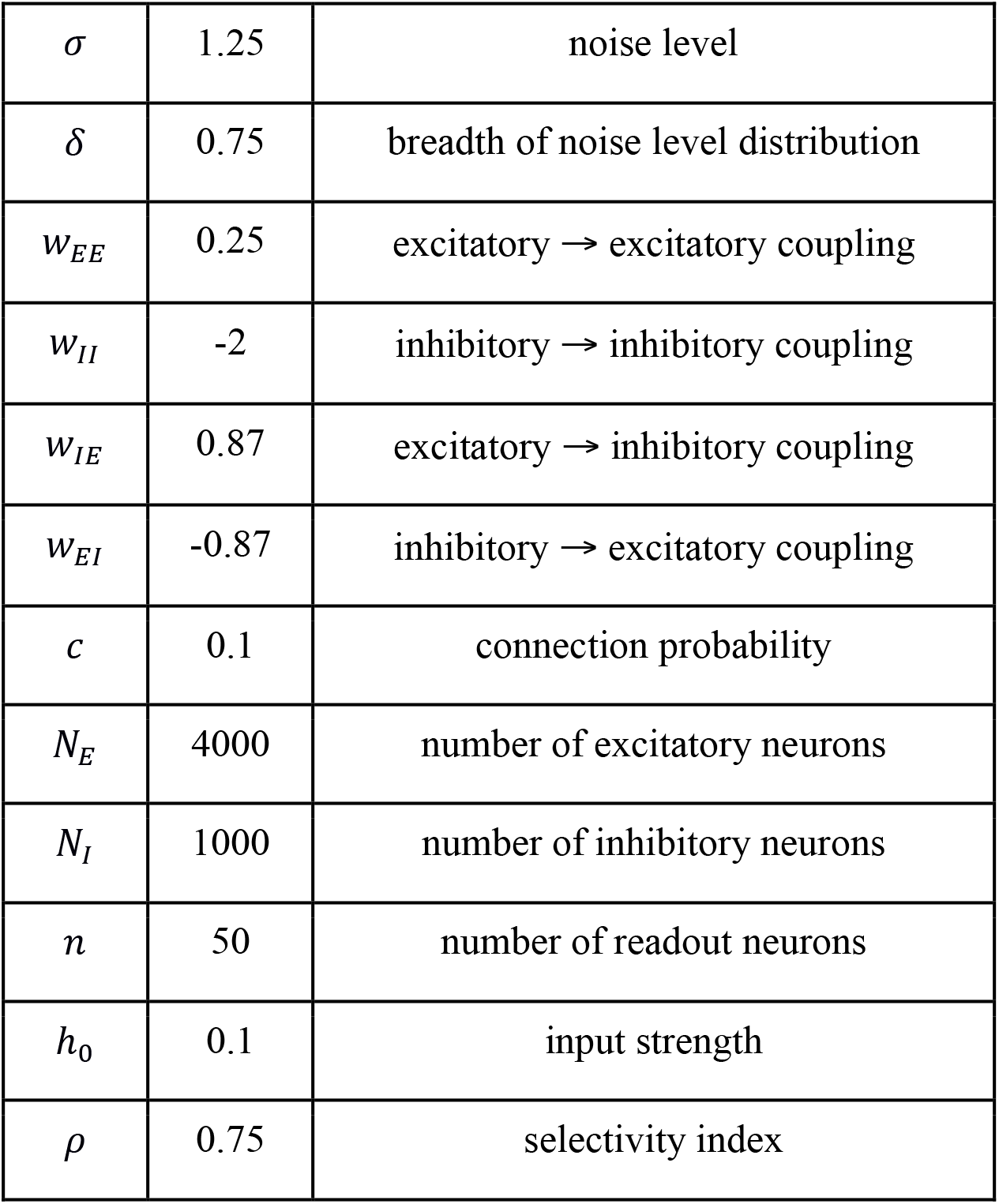
Parameters used in simulations

|  |  |  |
| --- | --- | --- |
| $\sigma$ | 1.25 | noise level |
| $\delta$ | 0.75 | breadth of noise level distribution |
| $w_{EE}$ | 0.25 | excitatory → excitatory coupling |
| $w_{II}$ | -2 | inhibitory → inhibitory coupling |
| $w_{IE}$ | 0.87 | excitatory → inhibitory coupling |
| $w_{EI}$ | -0.87 | inhibitory → excitatory coupling |
| $c$ | 0.1 | connection probability |
| $N_E$ | 4000 | number of excitatory neurons |
| $N_I$ | 1000 | number of inhibitory neurons |
| $n$ | 50 | number of readout neurons |
| $h_0$ | 0.1 | input strength |
| $\rho$ | 0.75 | selectivity index |

The goal is to determine the value of *s* (that is, determine whether **h**_1_ or **h**_2_ was present) given the activity of a subset of the neurons from either the excitatory or inhibitory populations. We’ll work in steady state, for which

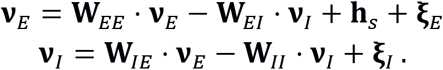

Solving for **ν**_*E*_ and **ν**_*I*_ yields

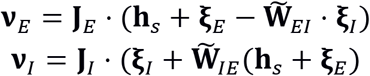

where

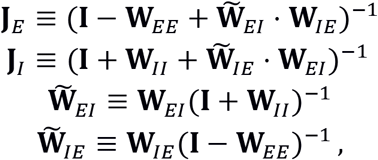

and **I** is the identity matrix. We are interested in the decoding accuracy of a sub-population of neurons. For that we’ll use a matrix **D**_*n*_ that picks out *n* components of whatever it’s operating on. So, for instance, **D**_*n*_ · **ν**_*E*_ is an *n*-dimensional vector with components equal to *n* of the components of **ν**_*E*_.

For a linear and Gaussian model such as ours, in which the covariance is independent of *s*, we need two quantities to compute the performance of an optimal decoder: the difference in the means of the subsampled populations when **h**_1_ versus **h**_2_ are present, and covariance matrix of the subsampled populations. The difference in means are given by

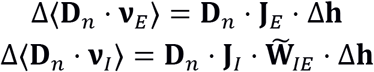

where Δ**h** is the difference between the two inputs,

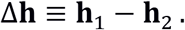

The covariances are given by

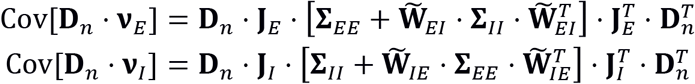

where *T* denotes transpose. Combining the mean and covariance gives us the signal to noise ratio,

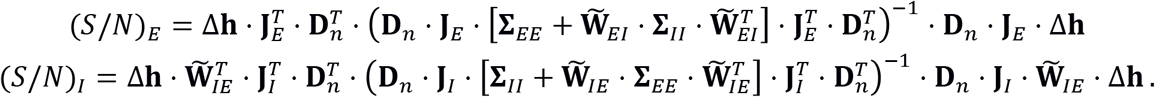

The performance of an optimal decoder is then given by

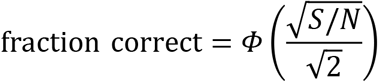

where *Φ* is the cumulative normal function. All of our analysis is based on this expression. Differences in fraction correct depend only on differences in the connectivity matrices, which we describe next.

#### Connectivity matrices

We consider three connectivity structures: completely non-selective, signal-selective, and signal-to-noise selective (corresponding to Figures 5A, 5B and 5C, respectively). In all cases the connectivity is sparse (the connection probability between any two neurons is 0.1). What differs is the connection strength when neurons are connected. We describe below how the connection strength is chosen for our three connectivity structures.

##### Non-selective

The connectivity matrices have the especially simple form

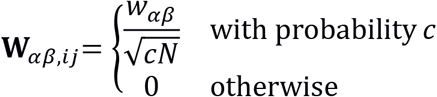

where *α, β* ∈ {*E, I*}, *N*(≡ *N_E_* + *N_I_*) is the total number of neurons, and *w_αβ_* are parameters (see Table 1).

##### Signal-selective

We divide the neurons into two sets of excitatory and inhibitory subpopulations, as in Figure 5B. The connection strengths are still given by the above expression, but now *α* and *β* acquire subscripts that specify which population they are in: *α, β* ∈ [*E*_1_, *E*_2_, *I*_1_ *I*_2_}, with *E_t_* and *I*_1_ referring to population 1 and *E*_2_ and *I*_2_ to population 2. The within-population connection strengths are the same as for the non-selective population (*w_α_i_β_i__* = *w_αβ_, i* = 1,2), but the across-population connection strengths are smaller by a factor of *ρ*,

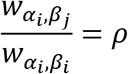

for *i* = 1 and *j* = 2 or vice-versa. The value of *ρ* determines how selective the sub-populations are: *ρ* = 0 corresponds to completely selective sub-populations while *ρ* = 1 corresponds to the completely non-selective case.

##### SNR-selective

We choose the connectivity as in the non-selective case, and then change synaptic strength so that the inhibitory neurons receive stronger connections from the excitatory neurons with high signal to noise ratios. To do that, we first rank excitatory units in order of ascending signal to noise ratio (by using **D**_1_ in the expression for (*S/N*)_*E*_ in the previous section). We then make the substitution

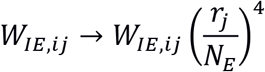

where *r_j_* is the rank of excitatory *j* in the order of ascending signal to noise ratio and, recall, *N_E_* is the number of excitatory neurons. This downweights projections from low signal to noise ratio excitatory neurons and upweights connections from high signal to noise ratio neurons. Finally, all elements are scaled to ensure that the average connection strength from the excitatory to the inhibitory network is the same as before the substitution.

#### Simulation details

The simulation parameters are given in Table 1. In addition, there are a number of relevant details, the most important of which is related to the input, **h**_*s*_. As mentioned in the previous section, about half the elements of **h**_*s*_ are *h*_0_ for the rightward choice and – *h*_0_ for the leftward choice, and the rest are *h*_0_ for the leftward choice – *h*_0_ for the rightward choice. This is strictly true for the completely non-selective and signal to noise selective connectivity; for the signal selective connectivity, we use **h**_*s,i*_ = *h*_0_ for the rightward choice and – *h*_0_ for the leftward choice when excitatory neuron *i* is in population 1, and **h**_*s,i*_ = *h*_0_ for the leftward choice and –*h*_0_ for the rightward choice when excitatory neuron *i* is in population 2. In either case, however, this introduces a stochastic element: for the completely non-selective and signal to noise selective connectivities, there is randomness in both the input and the circuit; for the signal selective connectivity, there is randomness in the circuit. In the former case, we can eliminate the randomness in the connectivity by averaging over the input, as follows.

Because the components of Δ**h** are independent, we have

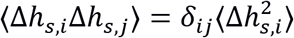

where *δ_ij_* is the Kronecker delta (*δ_ij_* = 1 if *i* = *j* and zero otherwise). Because Δ*h_s,i_* is either +*h*_0_ or −*h*_0_, we have

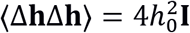

where **I** is the identity matrix. Thus, when we average the signal to noise ratios over Δ**h**, the expressions simplify slightly,

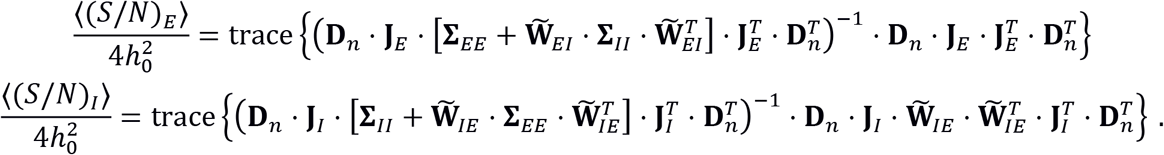

To avoid having to numerically average over input, we used these expressions when computing decoding accuracy for the completely non-selective and signal to noise selective connectivity. That left us with some randomness associated with the networks (as connectivity is chosen randomly), but that turned out to produce only small fluctuations, so each data point in Figures 5A and 5C was from a single network. For the signal selective connectivity (Figure 5B), the network realization turned out to matter, so we averaged over 25 networks, and for each of them we did a further averaging over 100 random picks of the 50 neurons from which we decoded.

In Figure 5, the x-axis is the ratio of the average connection strength from excitatory to inhibitory neurons to the average connection strength from inhibitory to excitatory neurons. This was chosen because it turned out to be the connectivity parameter with the largest effect on decoding accuracy. That in turn is because it turns out to be equivalent to the input noise to the inhibitory population. To see why, make the substitution

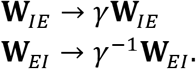

By letting **ν**_*I*_ → *γ***ν**_*I*_, we see that this is formally equivalent to letting **ξ**_*I*_ → *γ*^−1^**ξ**_*I*_, which in turn corresponds to letting **Σ**_*II*_ → *γ*^−2^**Σ**_*II*_. Thus the x-axis in Figure 5 can be thought of as the axis of decreasing input noise to the inhibitory neurons.

### Data and code availability

All the data used in the paper are publicly available on CSHL repository: http://repository.cshl.edu/36980/. Further, all the data is converted into the NWB format (Neurodata Without Boarders (Teeters et al., 2015; Ruebel et al., 2019), and is available on CSHL repository: https://dx.doi.org/10.14224/1.37693

Code for data processing and analysis is publicly available on github: https://github.com/farznaj/imaging_decisionMaking_exc_inh

Code for converting data to NWB format is also available on github: https://github.com/vathes/najafi-2018-nwb

## Author Contributions

Conceptualization and Writing: FN and AKC. Experiments and Analysis: FN. Decoding methodology and common-slope regression model: GFE, JPC and FN. Circuit modeling: RC and PEL. Spike-inference methodology: EAP. Funding Acquisition, Resources and Supervision: AKC.

## Acknowledgements

We thank Hien Nguyen for help with training mice, Matt Kaufman, Kachi Odoemene, Fred Marbach for technical assistance and thoughtful conversations. We thank Andrea Giovannucci for help with ROI inclusion criteria #5. We thank Ashley Juavinett, Simon Musall, and Sashank Pisupati for helpful discussions and feedback on early versions of the manuscript. We thank Thinh Nguyen, Dimitri Yatsenko and Edgar Walker for help with data conversion into the NWB format. The work was supported by the Simons Collaboration on the Global Brain, ONR MURI, the Klingenstein-Simons Foundation, the Pew Charitable Trust and the Gatsby Charitable Foundation.

## Supplemental Figures

**Figure S1.**
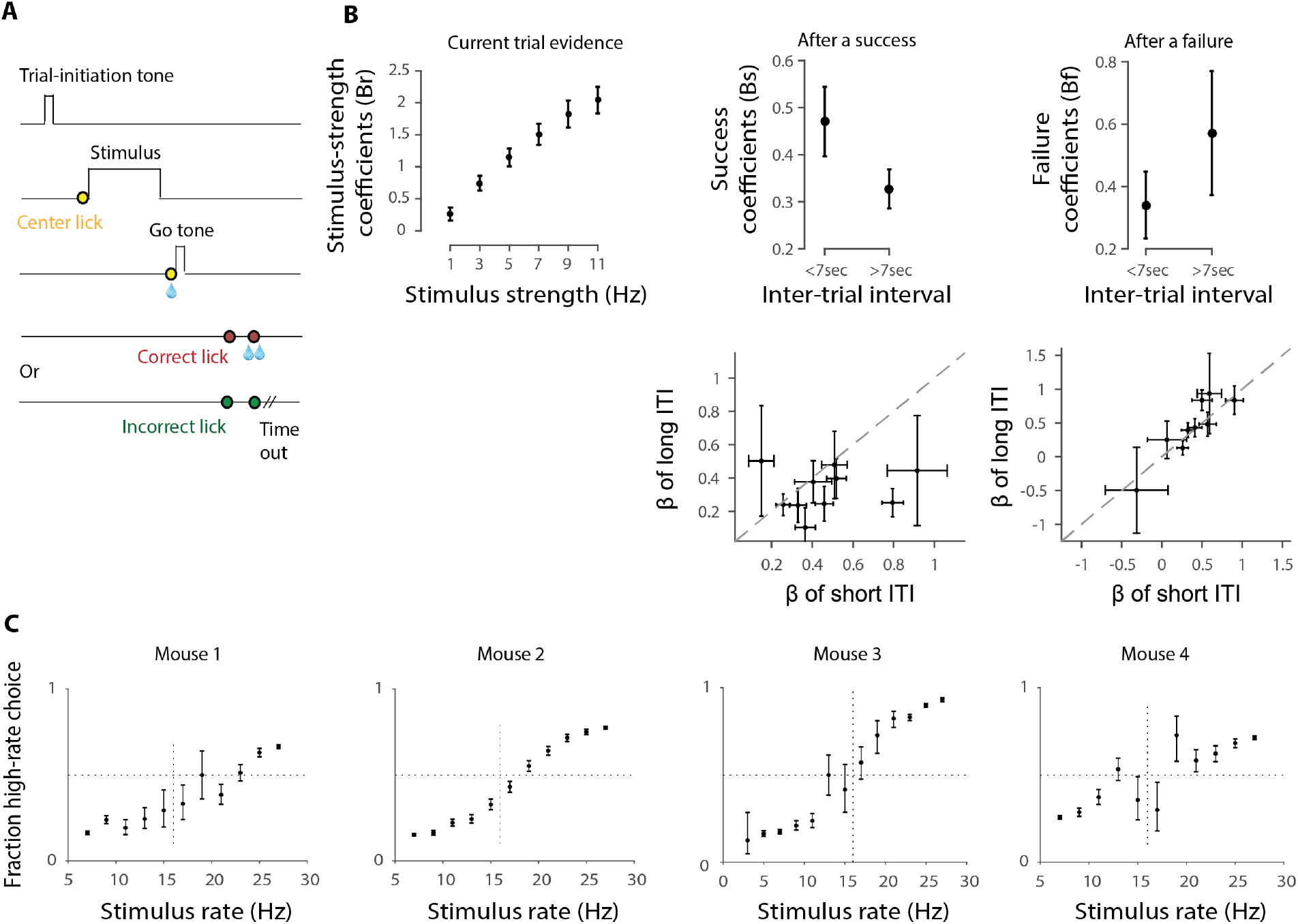
Related to Figure 1. Perceptual decisions about stimulus rate reflect current evidence, previous trial’s outcome, and the time passed since the previous trial. A, Trial structure. In each trial, first a brief tone is presented to indicate to the animal to initiate the trial (“trial-initiation tone”). Once the animal licks to the center waterspout (row 2: yellow circle), the stimulus is presented for 1 sec. At the end of the stimulus, the animal is required to lick again in the center (row 3: yellow circle). This will result in: 1) a small water reward in the center, 2) a “go tone” that indicates to the animal to make its choice. If the animal licks to the correct side (row 4, 1^st^ red circle), and confirms this lick (row 4, 2^nd^ red circle), it will receive water as a reward. If the animal licks to the wrong side (last row, 1^st^ green circle), and confirms this lick (last row, 2^nd^ green circle), there will be a time-out, i.e. longer time before the next trial can start. **B**, A logistic regression model was used to assess the extent to which the animal’s choice depends on stimulus strength (how far the stimulus rate is from the category boundary at 16Hz), previous choice outcome, and the time interval since the previous choice. Stimulus strength was divided into 6 bins (**left**); previous success was divided into 2 bins: success after a long ITI and success after a short ITI (middle); previous failure was also divided into 2 bins: failure after a long ITI and failure after a short ITI (right). Plots in top row show *β* averaged across animals (same 10 animals as in Figure 1B). Error bars: S.E.M across subjects. **Top Left:** strength of the sensory evidence affects the animal’s choices: the stronger the evidence, the higher the impact. Top **Middle:** Success of a previous trial also affects animal’s decision; the effect is stronger when the previous success occurs after a short ITI (<7sec). **Top Right:** Same but for previous incorrect trials; the effect of ITI after a failure was not significant. Plots in **bottom** row show success (left) and failure (right) *β* for individual mice. Error bars: S.E.M returned from glmfit.m in Matlab. **C**, Behavioral performance of the four mice in which we imaged excitatory and inhibitory activity during decision-making. In mice 1, 2, and 4, imaging was performed throughout learning by tracking the same group of neurons. Plots reflect data from all sessions. Errors bars: Wilson Binomial Confidence Interval.

**Figure S2.**
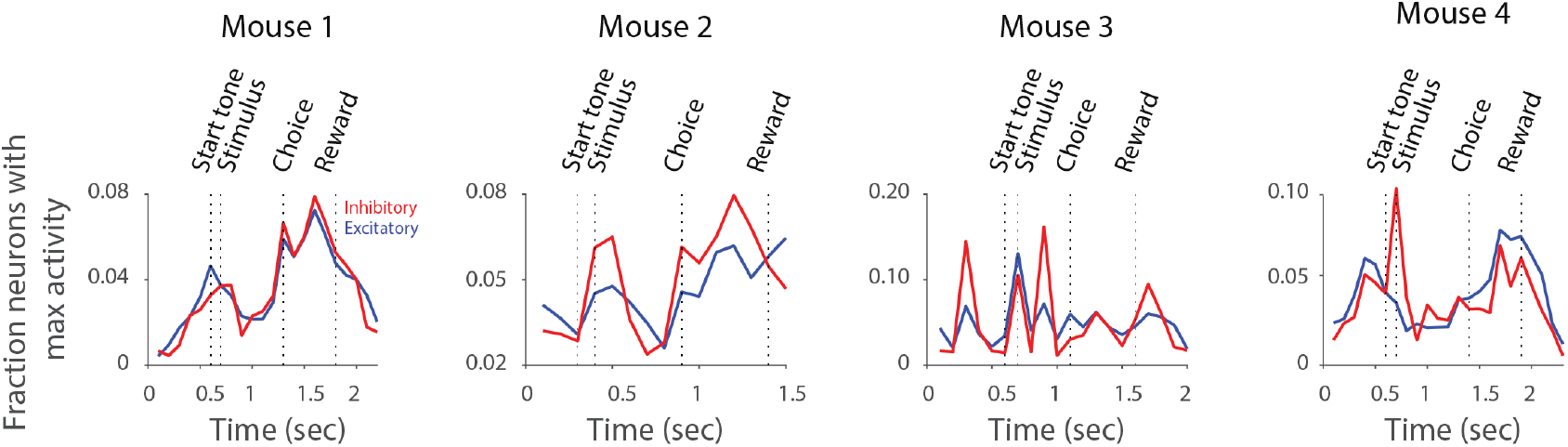
Related to Figure 1. Excitatory and inhibitory neurons have similar temporal dynamics. For each session, the fraction of neurons with peak activity in each 100ms time window was computed. This quantity is an estimate of the temporal-epoch tuning of neurons. Curves show mean across sessions, for excitatory (blue) and inhibitory (red) neurons, for each mouse. Similar to Figure 1E, traces were aligned for each trial event (start tone, stimulus, choice, reward), and then concatenated (see Figure 1E, legend).

**Figure S3.**
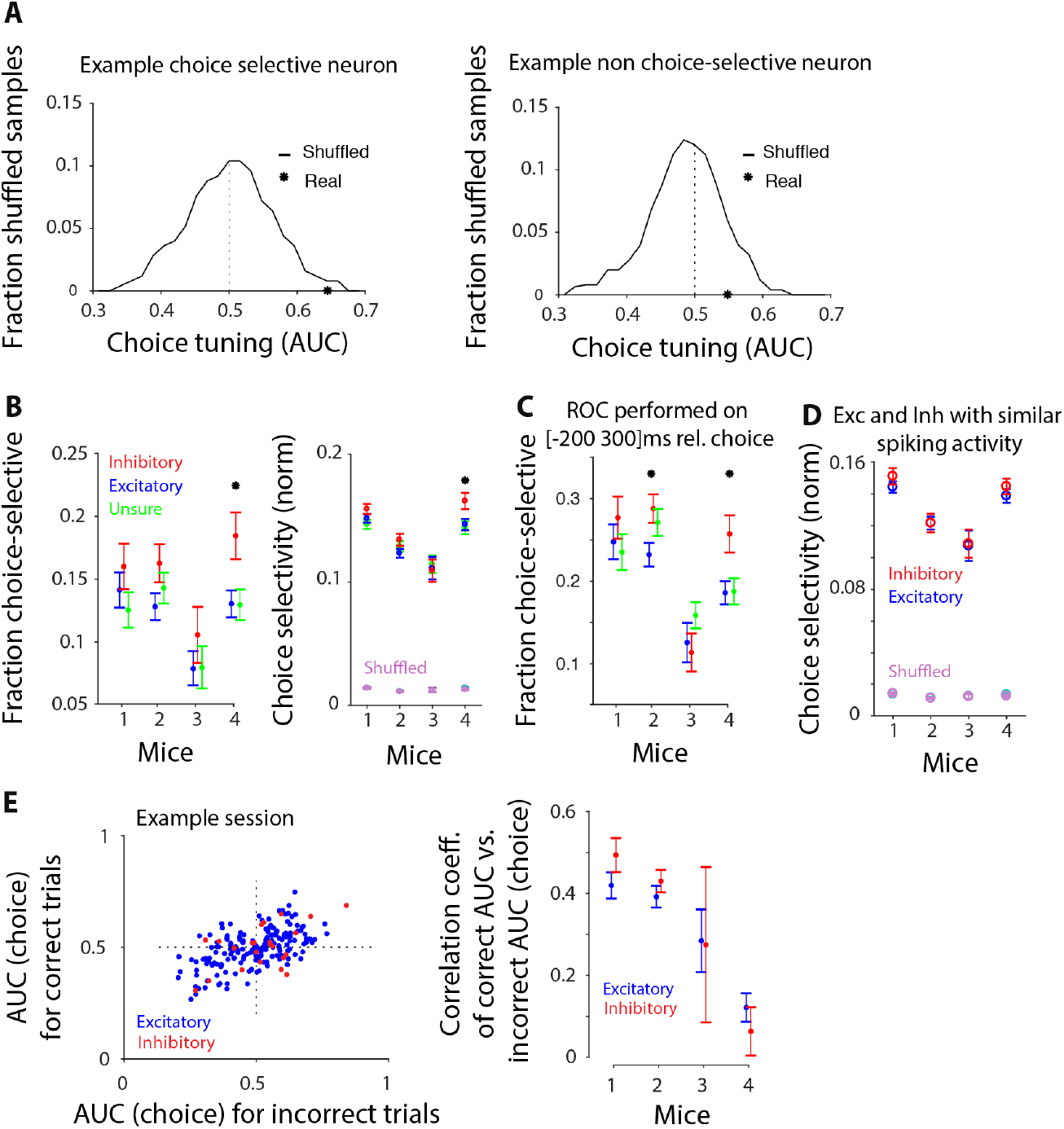
Related to Figure 2. Single neuron measures reveal similar choice selectivity in excitatory and inhibitory neurons. **A**, Example neurons to illustrate the method for assessing significant choice selectivity in individual neurons. In both panels, the solid line shows the distribution of values for the area under the ROC curve (AUC) generated by 50 different trial shuffles in which trials were randomly assigned to a left vs. right choice. Star indicates the actual AUC value of the neuron. Significance was assessed from the probability of occurrence of the actual AUC value in the shuffle distribution. When probabilities were <0.05, neurons were considered choice selective. Only the neuron on the left has significant choice selectivity. **B**, Fraction (left) and magnitude (right) of choice selectivity are shown for the unsure neurons (i.e. neurons classified as neither excitatory nor inhibitory; green), as well as excitatory (blue) and inhibitory (red) neurons. Data for each mouse show mean +/-standard error across sessions. **C**, Fraction of choice-selective neurons based on ROC analysis on [-200 300]ms relative to the choice. Fraction selective neurons at this time window (median across mice): excitatory: 21%; inhibitory: 27%, resulting in approximately 11 inhibitory and 69 excitatory neurons with significant choice selectivity per session. There is a considerable increase in the fraction of selective neurons when using this time window rather than 0-97ms window (see Figure 2C for comparison). **D**, ROC analysis restricted to those excitatory and inhibitory neurons that had the same spiking activity. Choice selectivity is still similar between the two cell types. Note that the significant difference observed for mouse 4 in Figure 2C is absent after controlling for the difference in spiking activity of inhibitory and excitatory neurons. Mean +/- standard error across sessions. **E, left:** Choice selectivity was computed on correct trials (vertical axis) as well as error trials (horizontal axis), and was correlated between the two conditions. Data is from a single session, each point shows an individual neuron whose cell type is indicated by its color. The positive correlation indicates that choice selectivity was overall similar on correct and error trials (Pearsons’ correlation coefficient, excitatory neurons: r=0.58; p<0.001; inhibitory neurons: r=0.55, p=0.007). The small number of points in quadrants 2 and 4 indicate less frequent neurons that showed opposite selectivity on correct vs. error trials. **Right**, Summary of correlation coefficient of AUC on correct trials vs. AUC on incorrect trials, mean across sessions for each animal. Error bars: S.E.M. across sessions. The weaker correlation in mouse 4 indicates that this animal had a mixture of cells selective for the stimulus and cells selective for the choice. Note that although the center of the imaging window was identical in all animals, the imaging location within the window of this animal was slightly posterior to the others. The enrichment of cells selective for the stimulus, in this mouse compared to other mice, may reflect that the region we imaged in mouse 4 was closer to primary visual cortex.

**Figure S4.**
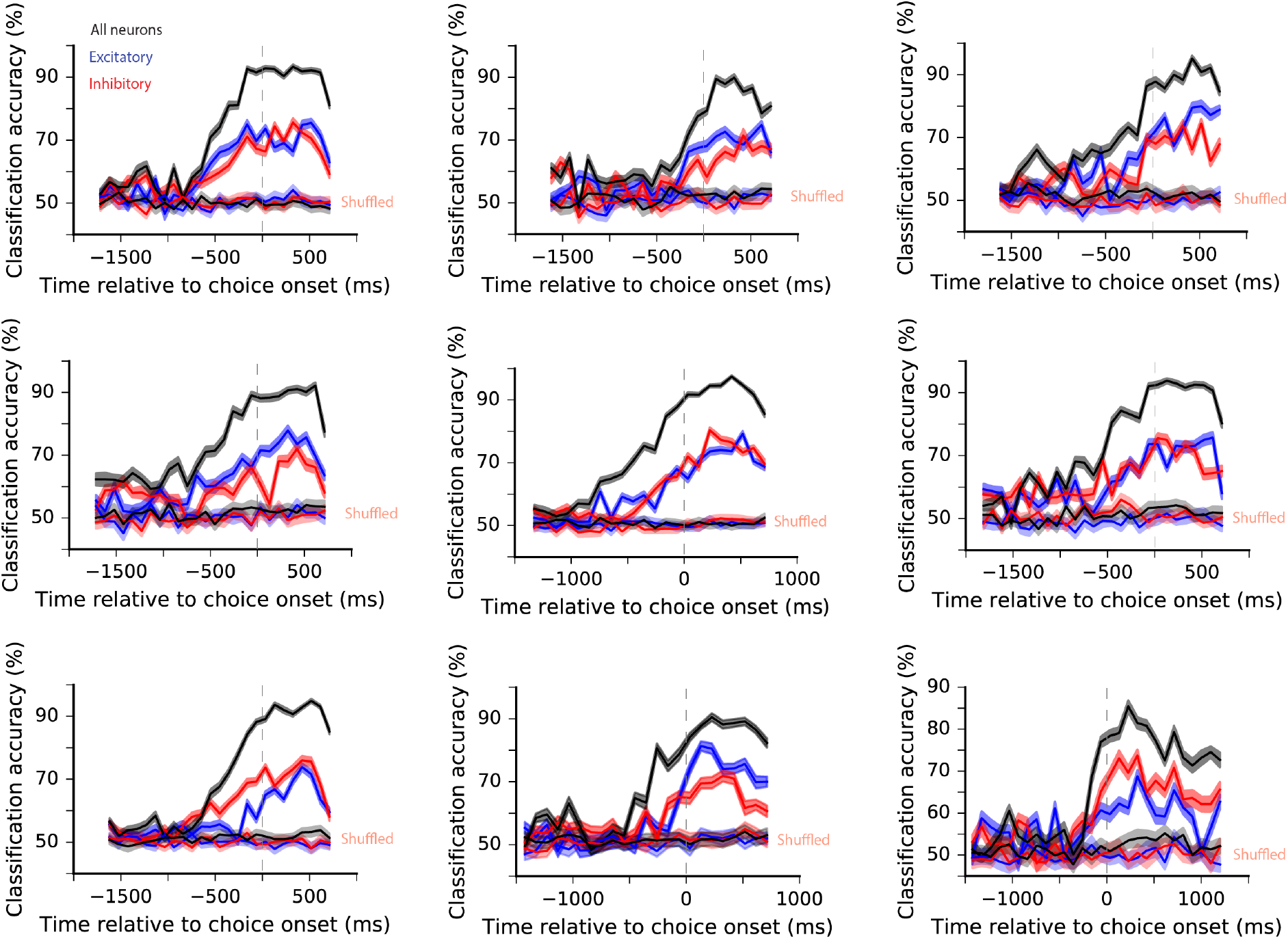
Related to Figure 3. Population activity is highly selective for the animal’s choice; excitatory and inhibitory neurons are similarly selective. Classification accuracy of the choice decoder at each moment in the trial for 9 additional example sessions. Dashed lines: choice onset. Black: all neurons included in the decoder; blue: subsampled excitatory neurons; red: inhibitory neurons; dim colors: shuffled control. In most sessions, inhibitory and subsampled excitatory populations have comparable classification accuracy.

**Figure S5.**
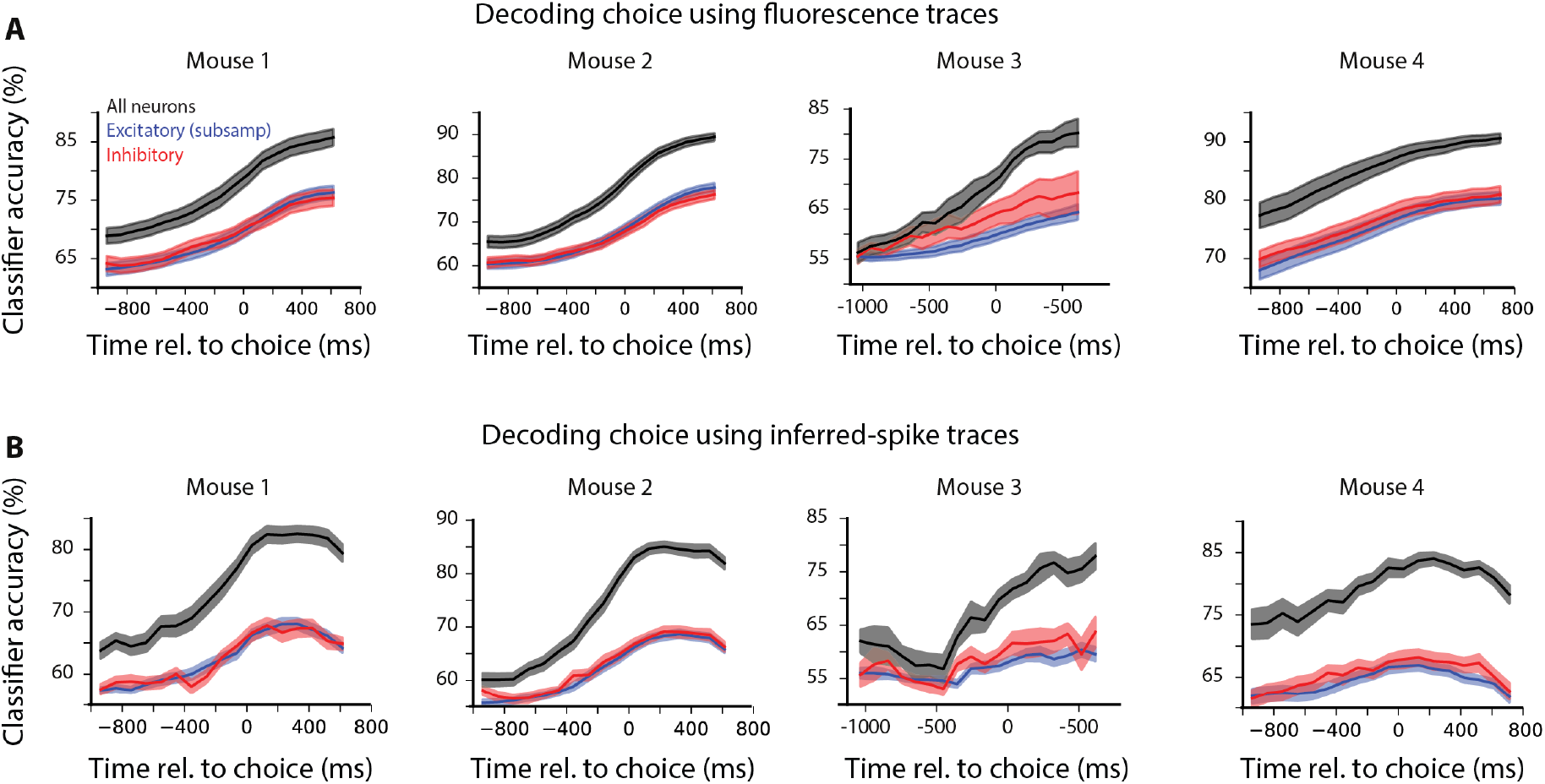
Related to Figure 3. Classification accuracy is similar for excitatory and inhibitory populations, whether the choice decoder is trained/tested on fluorescence traces or on inferred spikes. SVM classifiers were trained to decode choice from the population activity of all neurons (black), inhibitory neurons (red), or subsampled excitatory neurons (blue). In **(A)** fluorescence traces (Figure 1D middle) were used, and in **(B)** inferred spikes (Figure 1D right) were used. In both cases, decoder accuracy is similarly high for excitatory and inhibitory neurons.

**Figure S6.**
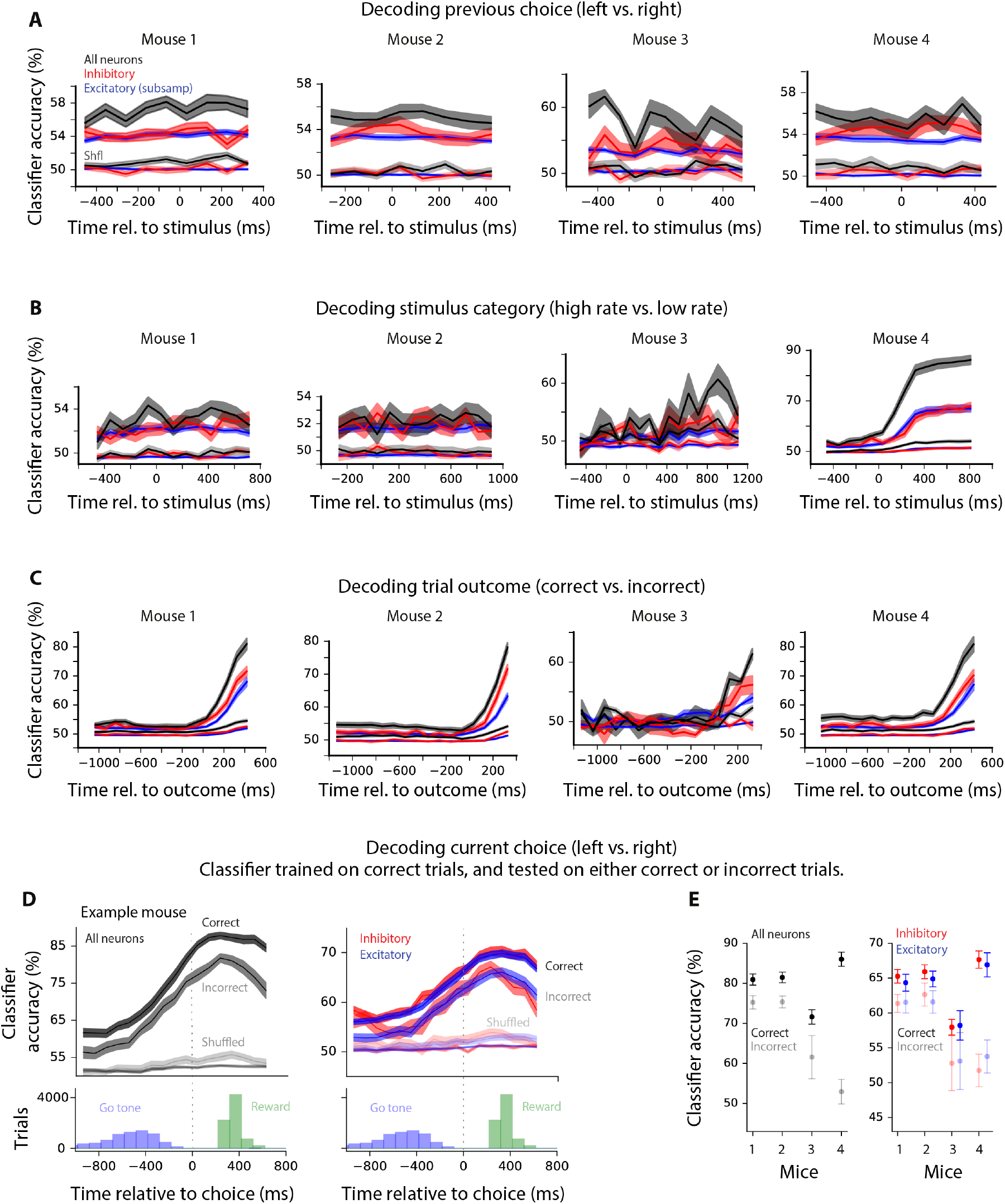
Related to Figure 3. Population activity is strongly selective for the trial outcome, and to a lesser degree to the stimulus category and previous choice. **A**, SVM classifier trained to decode previous choice from the activity of all neurons (black), inhibitory neurons (red), or subsampled excitatory (blue) neurons. “shfl” indicates classifier accuracy trained using shuffled trial labels. Previous choice is reflected, though weakly, in the population activity of the current trial. **B**, SVM classifier trained to decode the stimulus category, i.e. whether the stimulus is high rate (above 16Hz) or low rate (below 16Hz). Except for mouse 4, in which the imaging location was slightly more posterior (see Figure S3E, legend), stimulus category is weakly reflected in the population activity. **C**, SVM classifier trained to decode the trial outcome (i.e. correct vs. incorrect). Classification accuracy gradually increases and reaches 80% (median across mice) approximately 400ms after the animal confirms his choice (Figure S1A). Inhibitory neurons showed slightly higher selectivity for the outcome. Unsaturated lines in B and C: performance on shuffled trials. **D**, SVM classifier trained on correct trials to decode choice and tested on correct as well as incorrect trials. Data from an example animal (48 sessions). **Top**: Classification accuracy of decoders trained on all neurons (left), subsampled excitatory neurons (right, blue trace), and inhibitory neurons (right, red trace). In all cases, classifiers were trained on correct trials; however they were tested either on correct (dark lines: “Correct”) or incorrect (dim lines: “Incorrect”) trials. Classification accuracy on incorrect trials was high; indicating that population activity primarily reflects the animal’s choice, yet it differs at least slightly for correct and incorrect trials. This reduction was similar for excitatory and inhibitory neurons (blue are red traces are overlapping in the right panel). Bottom: Across-trial distribution of go tones and reward delivery (See Fig. 3B bottom). Left and right panels are the same plots and are duplicated to facilitate alignment to each corresponding plot above. **E**, Summary across all mice for all neurons (left) and excitatory and inhibitory neurons separately (right). Classifier performance on correct (dark colors) and incorrect (dim colors) trials is shown. Mouse 4 had the largest difference in classification accuracy for correct vs. error trials. As with the single-neuron analysis (Figure S3E) and decoding of stimulus category (Figure S6B), this difference likely reflects that the imaging region was slightly posterior within the window for this animal. Importantly, for all mice, the change in classification accuracy was quite similar for excitatory and inhibitory neurons (right), indicating that both populations reflect choice vs. stimulus to a comparable degree.

**Figure S7.**
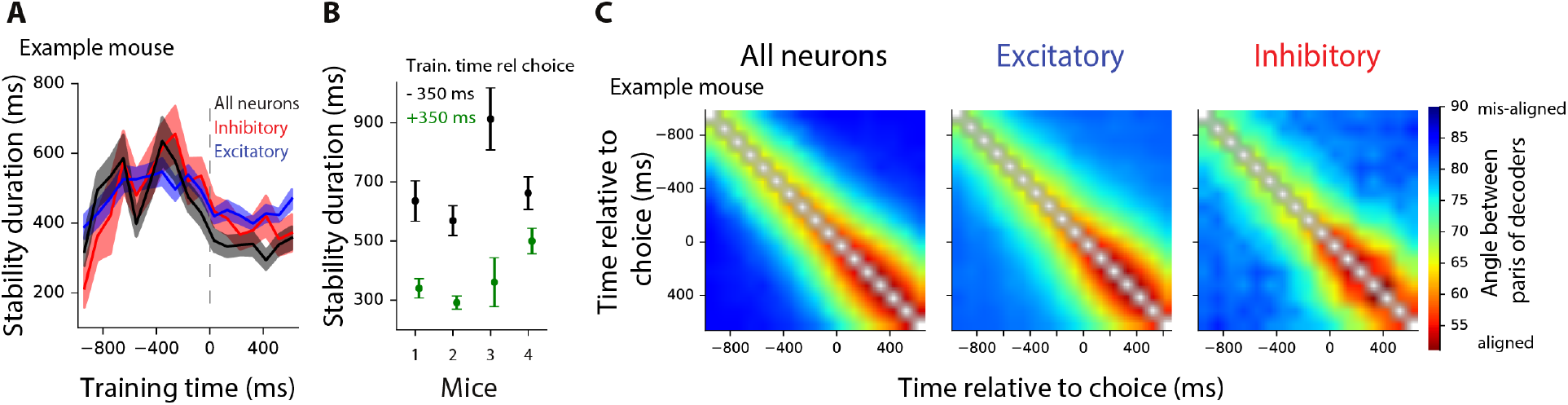
Related to Figure 4. Additional analyses provide more evidence for similar temporal stability of the choice decoder in excitatory and inhibitory populations. **A**, In an example mouse, population activity that predicts the animal’s choice is similarly stable for excitatory and inhibitory neurons during the course of a trial. The vertical axis shows the stability duration for decoders trained at different times during the trial. Stability duration is defined as the width of the testing window over which decoder accuracy does not statistically differ from that within the training window (red regions of Figure 4C) from that obtained by using the same training and testing times (diagonal of Figure 4A). Error bars: S.E.M. across sessions. Summary data for all mice at training time 0-97 ms before choice (dashed line) are shown in Figure 4D. **B**, Stability duration of the all-neuron decoder (black in panel A) is compared for decoders trained 350ms before the choice (black), and 350ms after the choice (green). Population stability was lower after the choice than before the choice. This may be due to additional events, e.g. reward delivery and repeated licking, which follow the choice. **C**, Another measurement of stability likewise suggests similar temporal stability for excitatory and inhibitory populations. Stability was assessed by measuring the angle between pairs of decoders trained at different time points in the trial. If a similar pattern of population activity represents choice from moment t_1_ to moment t_2_, the choice classifiers trained at these times will be aligned, i.e. the angle between the two classifiers will be small. The colors indicate the angle between pairs of decoders trained at different moments in the trial. Small angles (hot colors) indicate alignment of choice decoders; hence stable activity patterns, related to choice, across neurons. left: all neurons; middle: excitatory neurons (subsampled to match the number of inhibitory neurons); right: inhibitory neurons. As with our other method (Figure 4), the time course of stability was similar for excitatory and inhibitory neurons.

**Figure S8.**
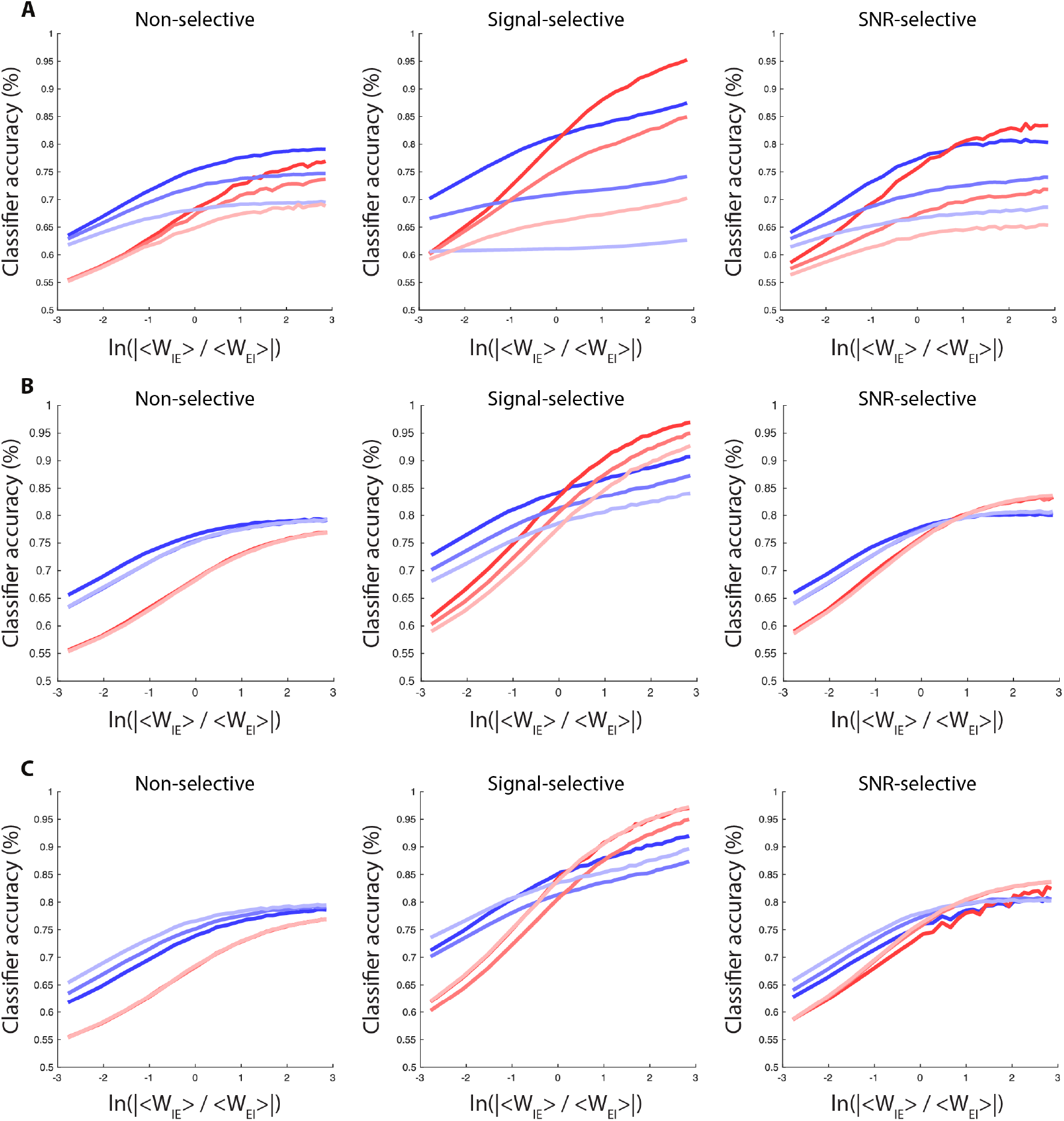
Related to Figure 5. Selective connectivity between excitatory and inhibitory neurons allows for matched classification accuracy in the two populations. Decoding accuracy versus three parameters. **A**, Differential correlations. **Σ**_*EE*_ → **Σ**_*EE*_ + *ϵΔ***h***Δ***h**. Dark to light hues: *ϵ* = 0, 17.78, 56.23. **B**, Excitatory to excitatory connections. Dark to light hues: *w_EE_* = 0.35, 0.3, 0.25 (default). **C**, Inhibitory to inhibitory connections. Dark to light hues: *w_II_* = −2.4, −2.0 (default), and −1.6.

**Figure S9.**
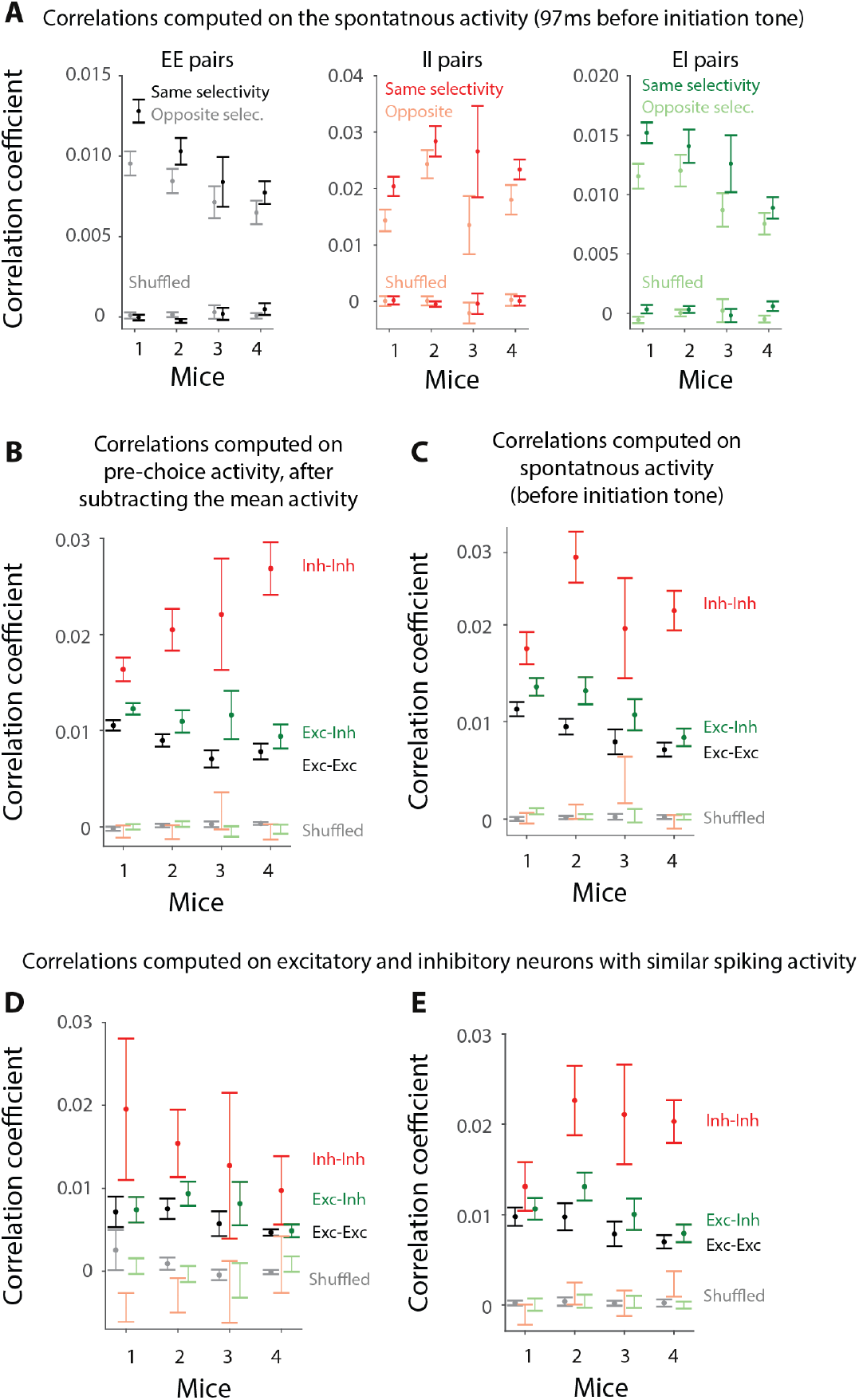
Related to Figure 6. Higher noise correlations between neurons with similar choice selectivity. Also, inhibitory neurons are more strongly correlated. **A**, Noise correlations between neurons with the same choice selectivity (dark colors) vs. those with opposite choice selectivity (dim colors), for pairs of excitatory neurons (left), pairs of inhibitory neurons (middle) or excitatory-inhibitory pairs (right). Signal correlations were not present because correlations were computed 0-97 ms before the trial initiation tone, when the stimulus is not present, and the activity is spontaneous. **B**, Noise correlations were much stronger for inhibitory-inhibitory pairs (red) than excitatory-excitatory pairs (black), and had intermediate values for excitatory-inhibitory pairs (green). Correlations are computed on 0-97ms before the choice after subtracting off the mean choice activity, hence removing the signal correlations. **C**, Same as B but for the time period 0-97 ms before the trial initiation tone (i.e. the spontaneous activity). **D,E**, same as in B,C, except correlations were computed only on those excitatory and inhibitory neurons with the same median spiking activity (Methods).

**Figure S10.**
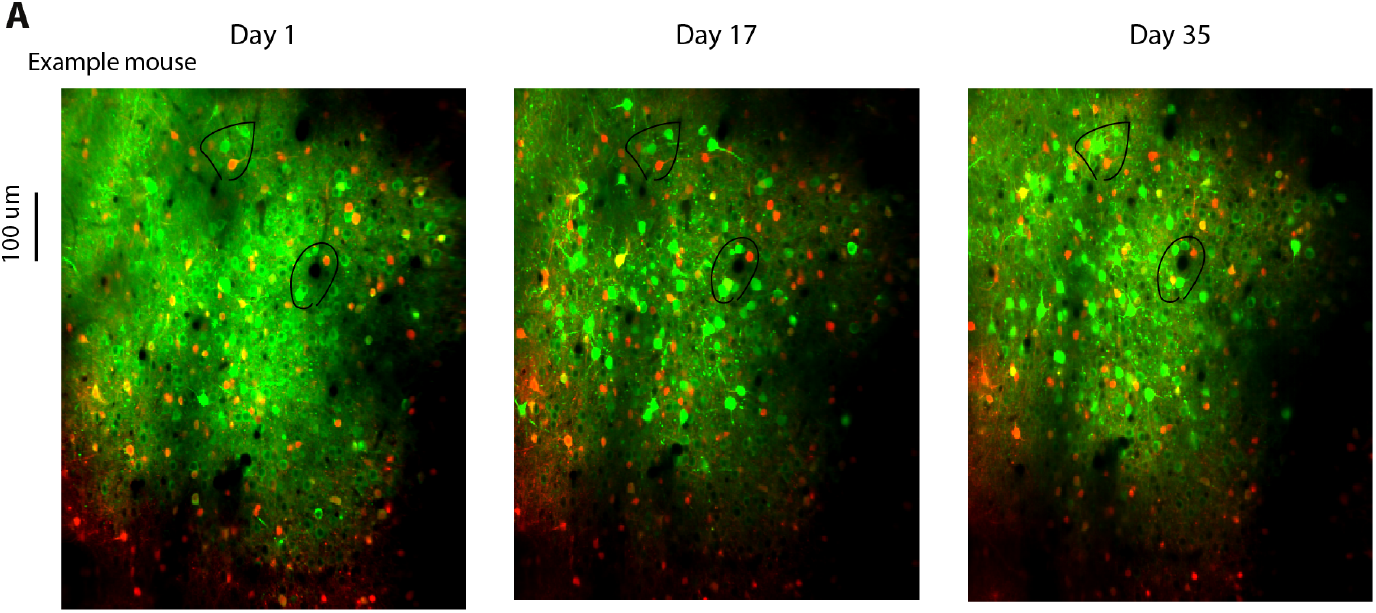
Related to Figure 8. The same field of view was imaged during learning. **A**, Field of view from three example sessions of a mouse: 1^st^ days of imaging (left), a middle imaging session (middle), and last day of imaging (right). Left to right panels span 60 days, out of which 35 days were experimental days. Black circles mark example areas that can be easily matched among the sessions. Each panel is an average image of all the frames imaged in the session. Green and red (bleedthrough corrected) images were merged.

**Figure S11.**
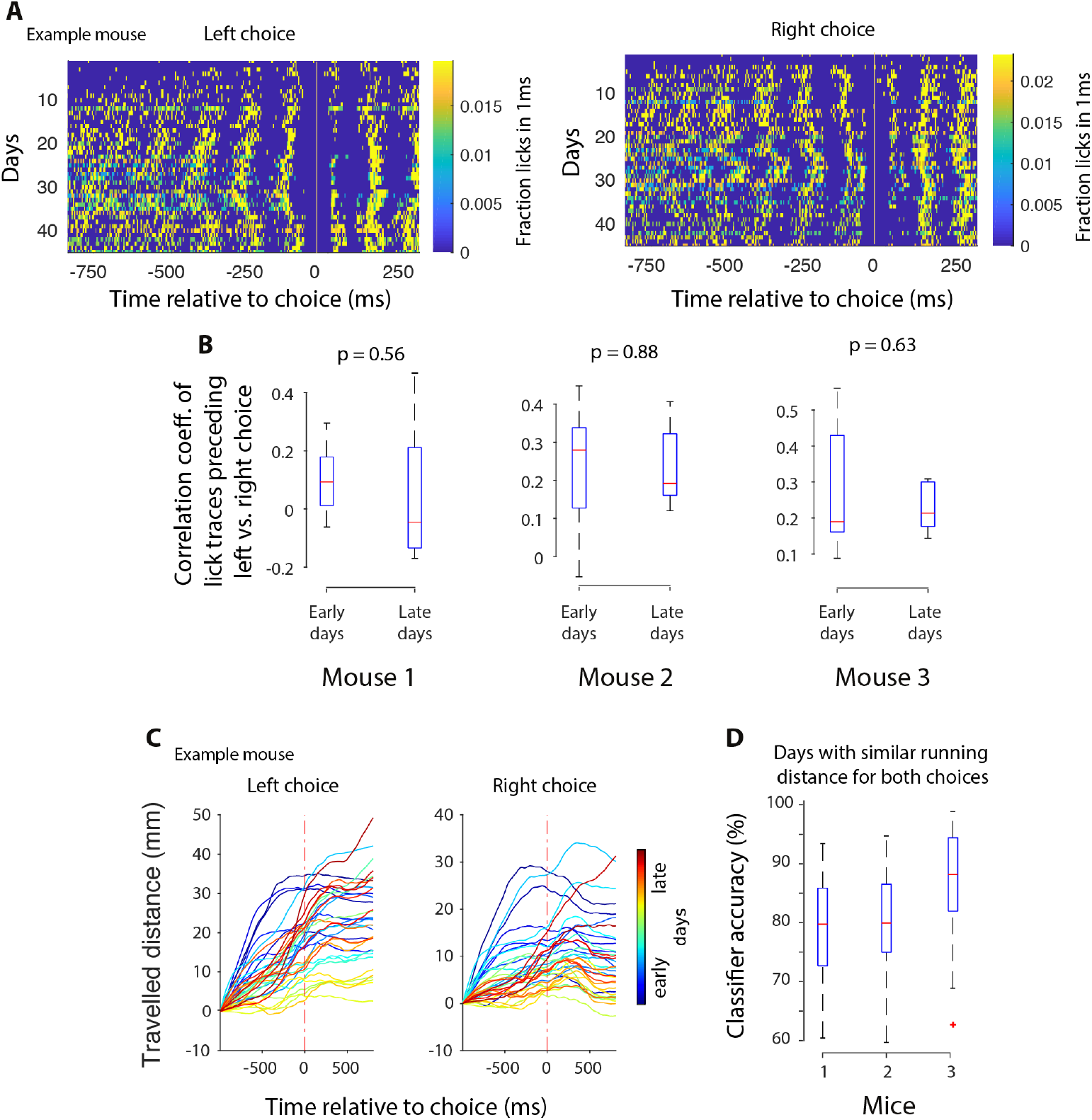
Related to Figure 8. Further analysis of learning-induced changes in the population activity: changes in licking and running movements are unlikely to account for improved classifier accuracy during learning. **A**, Licking was similar in advance of high rate vs. low rate choices, both early and late in training. Licks that occur before the choice (vertical line at 0) are to the center waterspout, and licks that occur after the choice are to the side waterspouts; example mouse. **B**, Each plot shows the Pearson’s correlation coefficient between licking patterns, to the center waterspout, preceding left and right choices, calculated 250ms before the choice. These correlations were typically similar for early vs. late training days, indicating that animal’s licking pattern preceding left vs. right choices did not change drastically over the course of learning. **C**, Distance that the animal travelled during the decision (as measured by the rotary encoder on the running wheel) was similar in advance of left vs. right choices; example mouse; each line represents a session (cold colors: early sessions; hot colors: late sessions). **D**, Classifier accuracy (0-97 ms before the choice) of the full population was high even when the analysis was restricted to sessions in which the distance travelled was not significantly different (t-test, P>0.05; time 0-97 ms before the choice) for left vs. right choices. This analysis was necessary because for some mice in some sessions, there were idiosyncratic differences between the distances travelled in advance of left vs. right choices. In (B) and (D), median (red horizontal line), inter-quartile range (blue box), and the entire range of data (dashed black lines) are shown. There is a single red ‘+’ at the bottom of mouse 3. What is the story there?

**Figure S12.**
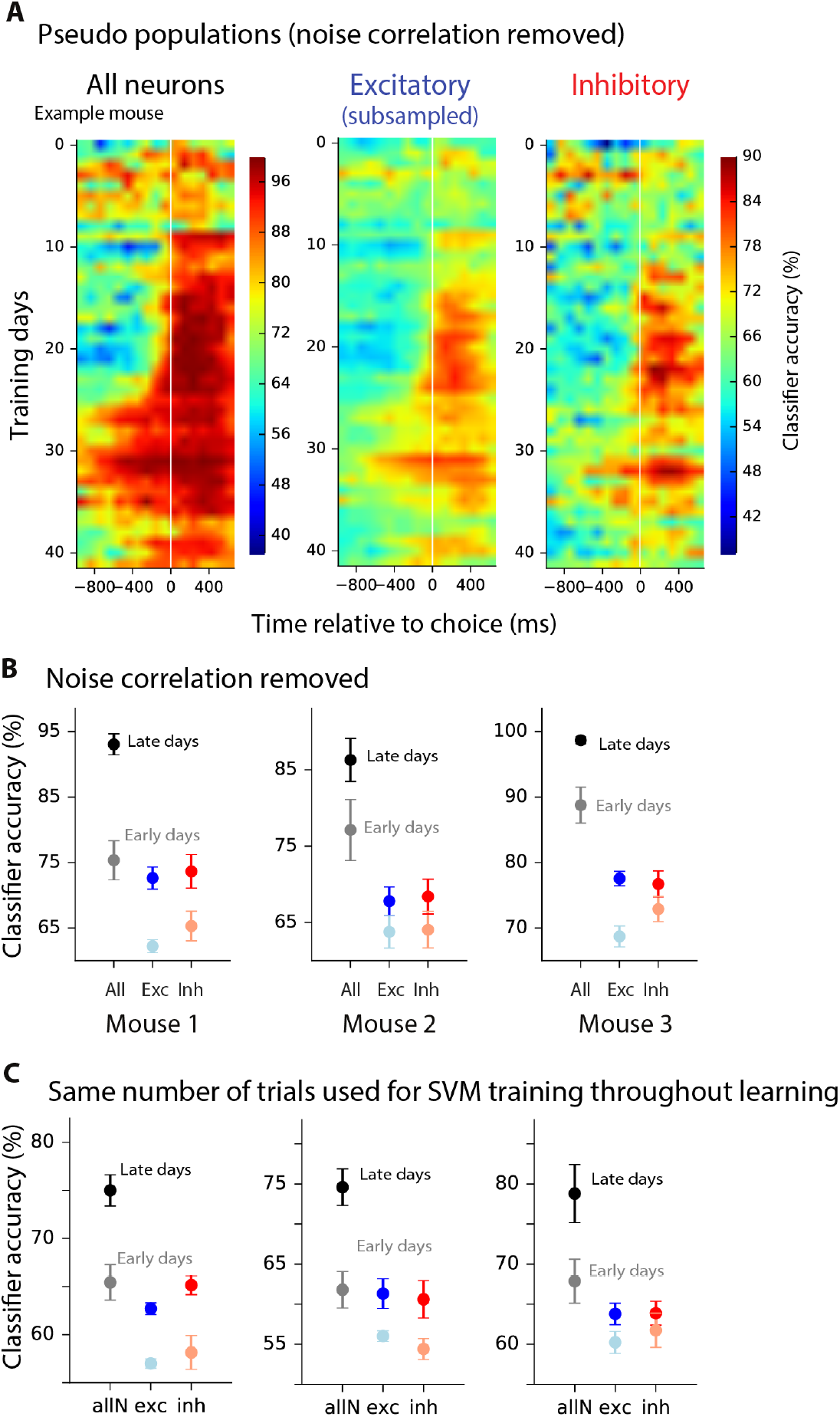
Related to Figure 8. Further analysis of learning-induced changes in the population activity: the reduction in noise correlations is insufficient to account for the improved classification accuracy during learning. Instead, the improvement can be explained by an increase in the fraction of significantly choice-selective neurons. **A**, Classification accuracy for each training session (average of cross-validation samples), for all neurons (left), subsampled excitatory (middle), and inhibitory neurons (right); example mouse. White vertical line: choice onset. This format is the same as Figure 8A, but here the noise correlations are removed by making pseudo populations (similar procedure as in Figure 7). **B**, Summary of each mouse, showing classification accuracy averaged across early (unsaturated colors) vs. late (saturated colors) training days, at 0-97ms before the choice. As in (A), data are based on pseudo-populations in which the noise correlations are removed. The learning-induced improvement in the classifier accuracy in pseudo populations indicates that reduced noise correlations (Figure 8F) cannot solely account for the enhanced classifier accuracy in the population during learning (Figure 8A). **C**, Equal trial numbers were used to train the choice classifier in every session to control for any effects of trial numbers on classifier accuracy. An increase in classifier accuracy is still observed as a result of learning. Classifiers were trained only on correct trials.

**Figure S13.**
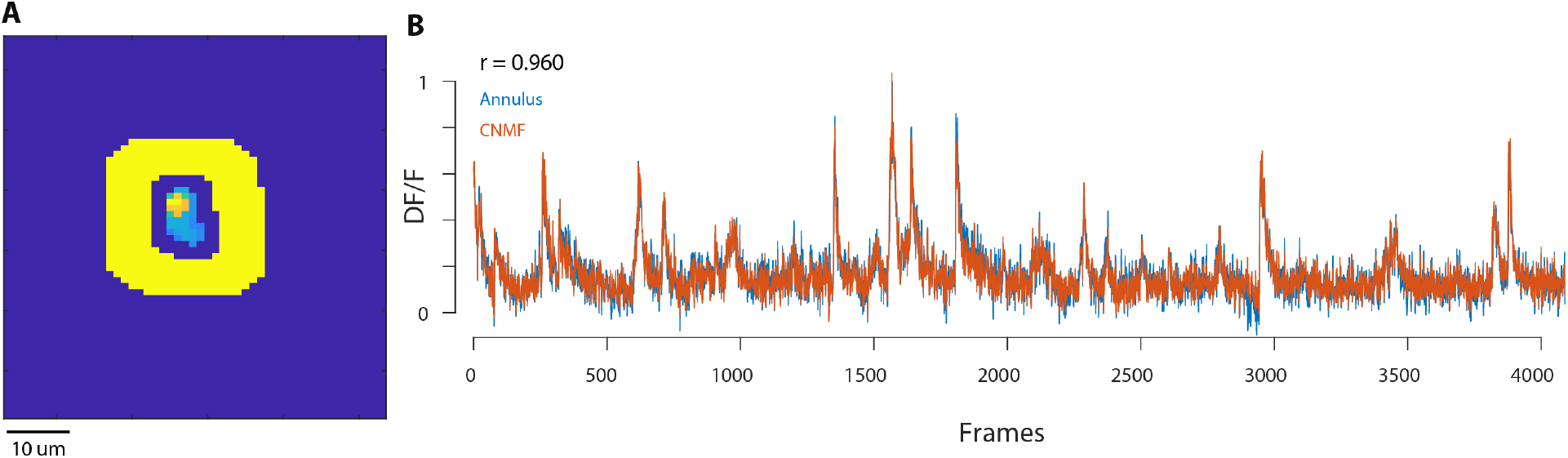
Related to Methods section “Neuropil Contamination removal”. Removing neuropil contamination with CNMF or manually using an annulus leads to the same results. **A**, An example spatial component in the FOV and its surrounding annulus (yellow). **B**, ΔF/F trace for the same component obtained by manually subtracting the neuropil activity averaged over the annulus region (blue trace) or by using the output of the CNMF processing pipeline (red trace). The two traces look nearly identical as also demonstrated by their high correlation coefficient (r = 0.96; the traces are not denoised). These results demonstrate the ability of the CNMF framework to properly capture neuropil contamination and remove it from the detected calcium traces.

## References

Akrami A, Kopec CD, Diamond ME, Brody CD (2018) Posterior parietal cortex represents sensory history and mediates its effects on behaviour. Nature 554:368–372.

Allen WE, Kauvar IV, Chen MZ, Richman EB, Yang SJ, Chan K, Gradinaru V, Deverman BE, Luo L, Deisseroth K (2017) Global Representations of Goal-Directed Behavior in Distinct Cell Types of Mouse Neocortex. Neuron 94:891–907 e896.

Atallah BV, Bruns W, Carandini M, Scanziani M (2012) Parvalbumin-expressing interneurons linearly transform cortical responses to visual stimuli. Neuron 73:159–170.

Averbeck BB, Lee D (2006) Effects of noise correlations on information encoding and decoding. J Neurophysiol 95:3633–3644.

Averbeck BB, Latham PE, Pouget A (2006) Neural correlations, population coding and computation. Nat Rev Neurosci 7:358–366.

Beaulieu C (1993) Numerical data on neocortical neurons in adult rat, with special reference to the GABA population. Brain Res 609:284–292.

Beck JM, Ma WJ, Kiani R, Hanks T, Churchland AK, Roitman J, Shadlen MN, Latham PE, Pouget A (2008) Probabilistic population codes for Bayesian decision making. Neuron 60:1142–1152.

Bock DD, Lee WC, Kerlin AM, Andermann ML, Hood G, Wetzel AW, Yurgenson S, Soucy ER, Kim HS, Reid RC (2011) Network anatomy and in vivo physiology of visual cortical neurons. Nature 471:177–182.

Bogacz R, Brown E, Moehlis J, Holmes P, Cohen JD (2006) The physics of optimal decision making: a formal analysis of models of performance in two-alternative forced-choice tasks. Psychol Rev 113:700–765.

Brunton BW, Botvinick MM, Brody CD (2013) Rats and humans can optimally accumulate evidence for decision-making. Science 340:95–98.

Busse L, Ayaz A, Dhruv NT, Katzner S, Saleem AB, Scholvinck ML, Zaharia AD, Carandini M (2011) The detection of visual contrast in the behaving mouse. J Neurosci 31:11351–11361.

Ch’ng YH, Reid RC (2010) Cellular imaging of visual cortex reveals the spatial and functional organization of spontaneous activity. Front Integr Neurosci 4.

Chen TW, Wardill TJ, Sun Y, Pulver SR, Renninger SL, Baohan A, Schreiter ER, Kerr RA, Orger MB, Jayaraman V, Looger LL, Svoboda K, Kim DS (2013) Ultrasensitive fluorescent proteins for imaging neuronal activity. Nature 499:295–300.

Constantinidis C, Goldman-Rakic PS (2002) Correlated discharges among putative pyramidal neurons and interneurons in the primate prefrontal cortex. J Neurophysiol 88:3487–3497.

Cossell L, Iacaruso MF, Muir DR, Houlton R, Sader EN, Ko H, Hofer SB, Mrsic-Flogel TD (2015) Functional organization of excitatory synaptic strength in primary visual cortex. Nature 518:399–403.

de Lima AD, Voigt T (1997) Identification of two distinct populations of gamma-aminobutyric acidergic neurons in cultures of the rat cerebral cortex. J Comp Neurol 388:526–540.

de Lima AD, Gieseler A, Voigt T (2009) Relationship between GABAergic interneurons migration and early neocortical network activity. Dev Neurobiol 69:105–123.

Deneve S, Latham PE, Pouget A (1999) Reading population codes: a neural implementation of ideal observers. Nat Neurosci 2:740–745.

Driscoll LN, Pettit NL, Minderer M, Chettih SN, Harvey CD (2017) Dynamic Reorganization of Neuronal Activity Patterns in Parietal Cortex. Cell 170:986–999 e916.

Ego-Stengel V, Wilson MA (2007) Spatial selectivity and theta phase precession in CA1 interneurons. Hippocampus 17:161–174.

Elsayed GF, Lara AH, Kaufman MT, Churchland MM, Cunningham JP (2016) Reorganization between preparatory and movement population responses in motor cortex. Nat Commun 7:13239.

Francis NA, Winkowski DE, Sheikhattar A, Armengol K, Babadi B, Kanold PO (2018) Small Networks Encode Decision-Making in Primary Auditory Cortex. Neuron 97:885–897 e886.

Freedman DJ, Assad JA (2006) Experience-dependent representation of visual categories in parietal cortex. Nature 443:85–88.

Funamizu A, Kuhn B, Doya K (2016) Neural substrate of dynamic Bayesian inference in the cerebral cortex. Nat Neurosci 19:1682–1689.

Gabbott PL, Dickie BG, Vaid RR, Headlam AJ, Bacon SJ (1997) Local-circuit neurones in the medial prefrontal cortex (areas 25, 32 and 24b) in the rat: morphology and quantitative distribution. J Comp Neurol 377:465–499.

Galarreta M, Hestrin S (1999) A network of fast-spiking cells in the neocortex connected by electrical synapses. Nature 402:72–75.

Giovannucci A, Friedrich J, Gunn P, Kalfon J, Koay SA, Taxidis J, Najafi F, Gauthier JL, Zhou P, Tank DW, Chklovskii DB, Pnevmatikakis EA (2018) CaImAn: An open source tool for scalable Calcium Imaging data Analysis. bioRxiv.

Giovannucci A, Friedrich J, Gunn P, Kalfon J, Brown BL, Koay SA, Taxidis J, Najafi F, Gauthier JL, Zhou P, Khakh BS, Tank DW, Chklovskii DB, Pnevmatikakis EA (2019) CaImAn an open source tool for scalable calcium imaging data analysis. Elife 8.

Goard MJ, Pho GN, Woodson J, Sur M (2016) Distinct roles of visual, parietal, and frontal motor cortices in memory-guided sensorimotor decisions. Elife 5.

Green DM, Swets JA (1966) Signal detection theory and psychophysics. New York: Wiley.

Gu Y, Liu S, Fetsch CR, Yang Y, Fok S, Sunkara A, DeAngelis GC, Angelaki DE (2011) Perceptual learning reduces interneuronal correlations in macaque visual cortex. Neuron 71:750–761.

Guizar-Sicairos M, Thurman ST, Fienup JR (2008) Efficient subpixel image registration algorithms. Opt Lett 33:156–158.

Harvey CD, Coen P, Tank DW (2012) Choice-specific sequences in parietal cortex during a virtual-navigation decision task. Nature 484:62–68.

Helmchen F, Tank DW (2019) A Single-Compartment Model of Calcium Dynamics in Nerve Terminals and Dendrites. Cold Spring Harb Protoc; doi:101101/pdbtop085910.

Hofer SB, Ko H, Pichler B, Vogelstein J, Ros H, Zeng H, Lein E, Lesica NA, Mrsic-Flogel TD (2011) Differential connectivity and response dynamics of excitatory and inhibitory neurons in visual cortex. Nat Neurosci 14:1045–1052.

Hofmann T, Scholkopf B, Smola AJ (2008) Kernel methods in machine learning. Ann Statist 36:1171–1220.

Hwang EJ, Dahlen JE, Mukundan M, Komiyama T (2017) History-based action selection bias in posterior parietal cortex. Nat Commun 8:1242.

Isaacson JS, Scanziani M (2011) How inhibition shapes cortical activity. Neuron 72:231–243.

Jeanne JM, Sharpee TO, Gentner TQ (2013) Associative learning enhances population coding by inverting interneuronal correlation patterns. Neuron 78:352–363.

Jouhanneau JS, Kremkow J, Poulet JFA (2018) Single synaptic inputs drive high-precision action potentials in parvalbumin expressing GABA-ergic cortical neurons in vivo. Nat Commun 9:1540.

Jouhanneau JS, Kremkow J, Dorrn AL, Poulet JF (2015) In Vivo Monosynaptic Excitatory Transmission between Layer 2 Cortical Pyramidal Neurons. Cell Rep 13:2098–2106.

Kamigaki T, Dan Y (2017) Delay activity of specific prefrontal interneuron subtypes modulates memory-guided behavior. Nat Neurosci 20:854–863.

Kerlin AM, Andermann ML, Berezovskii VK, Reid RC (2010) Broadly tuned response properties of diverse inhibitory neuron subtypes in mouse visual cortex. Neuron 67:85–8871.

Khan AG, Poort J, Chadwick A, Blot A, Sahani M, Mrsic-Flogel TD, Hofer SB (2018) Distinct learning-induced changes in stimulus selectivity and interactions of GABAergic interneuron classes in visual cortex. Nature Neuroscience.

Kim Y, Yang GR, Pradhan K, Venkataraju KU, Bota M, Garcia Del Molino LC, Fitzgerald G, Ram K, He M, Levine JM, Mitra P, Huang ZJ, Wang XJ, Osten P (2017) Brain-wide Maps Reveal Stereotyped Cell-Type-Based Cortical Architecture and Subcortical Sexual Dimorphism. Cell 171:456–469 e422.

Kimmel D, Elsayed GF, Cunningham JP, Rangel A, Newsome WT (2016) Encoding of value and choice as separable, dynamic neural dimensions in orbitofrontal cortex. Cosyne.

Ko H, Hofer SB, Pichler B, Buchanan KA, Sjostrom PJ, Mrsic-Flogel TD (2011) Functional specificity of local synaptic connections in neocortical networks. Nature 473:87–91.

Krishna VR, Alexander KR, Peachey NS (2002) Temporal properties of the mouse cone electroretinogram. J Neurophysiol 87:42–48.

Kwan AC, Dan Y (2012) Dissection of cortical microcircuits by single-neuron stimulation in vivo. Curr Biol 22:1459–1467.

Law CT, Gold JI (2008) Neural correlates of perceptual learning in a sensory-motor, but not a sensory, cortical area. Nat Neurosci 11:505–513.

Lee WC, Bonin V, Reed M, Graham BJ, Hood G, Glattfelder K, Reid RC (2016) Anatomy and function of an excitatory network in the visual cortex. Nature 532:370–374.

Lim S, Goldman MS (2013) Balanced cortical microcircuitry for maintaining information in working memory. Nat Neurosci 16:1306–1314.

Liu BH, Li P, Li YT, Sun YJ, Yanagawa Y, Obata K, Zhang LI, Tao HW (2009) Visual receptive field structure of cortical inhibitory neurons revealed by two-photon imaging guided recording. J Neurosci 29:10520–10532.

Lo CC, Wang XJ (2006) Cortico-basal ganglia circuit mechanism for a decision threshold in reaction time tasks. Nat Neurosci 9:956–963.

Lovett-Barron M, Kaifosh P, Kheirbek MA, Danielson N, Zaremba JD, Reardon TR, Turi GF, Hen R, Zemelman BV, Losonczy A (2014) Dendritic inhibition in the hippocampus supports fear learning. Science 343:857–863.

Ma WP, Liu BH, Li YT, Huang ZJ, Zhang LI, Tao HW (2010) Visual representations by cortical somatostatin inhibitory neurons--selective but with weak and delayed responses. J Neurosci 30:14371–14379.

Machado TA, Pnevmatikakis E, Paninski L, Jessell TM, Miri A (2015) Primacy of Flexor Locomotor Pattern Revealed by Ancestral Reversion of Motor Neuron Identity. Cell 162:338–350.

Machens CK, Romo R, Brody CD (2005) Flexible control of mutual inhibition: a neural model of two-interval discrimination. Science 307:1121–1124.

Madisen L, Zwingman TA, Sunkin SM, Oh SW, Zariwala HA, Gu H, Ng LL, Palmiter RD, Hawrylycz MJ, Jones AR, Lein ES, Zeng H (2010) A robust and high-throughput Cre reporting and characterization system for the whole mouse brain. Nat Neurosci 13:133–140.

Marbach F, Zador AM (2017) A self-initiated two-alternative forced choice paradigm for head-fixed mice. bioRxiv.

Maurer AP, Cowen SL, Burke SN, Barnes CA, McNaughton BL (2006) Phase precession in hippocampal interneurons showing strong functional coupling to individual pyramidal cells. J Neurosci 26:13485–13492.

Mi Y, Katkov M, Tsodyks M (2017) Synaptic Correlates of Working Memory Capacity. Neuron 93:323–330.

Moore AK, Wehr M (2013) Parvalbumin-expressing inhibitory interneurons in auditory cortex are well-tuned for frequency. J Neurosci 33:13713–13723.

Morcos AS, Harvey CD (2016) History-dependent variability in population dynamics during evidence accumulation in cortex. Nat Neurosci 19:1672–1681.

Moreno-Bote R, Beck J, Kanitscheider I, Pitkow X, Latham P, Pouget A (2014) Information-limiting correlations. Nat Neurosci 17:1410–1417.

Ni AM, Ruff DA, Alberts JJ, Symmonds J, Cohen MR (2018) Learning and attention reveal a general relationship between population activity and behavior. Science 359:463–465.

Niell CM, Stryker MP (2008) Highly selective receptive fields in mouse visual cortex. J Neurosci 28:7520–7536.

Odoemene O, Pisupati S, Nguyen H, Churchland AK (2017) Visual evidence accumulation guides decision-making in unrestrained mice. bioRxiv.

Packer AM, Yuste R (2011) Dense, unspecific connectivity of neocortical parvalbumin-positive interneurons: a canonical microcircuit for inhibition? J Neurosci 31:13260–13271.

Panzeri S, Schultz SR, Treves A, Rolls ET (1999) Correlations and the encoding of information in the nervous system. Proc Biol Sci 266:1001–1012.

Pfeffer CK, Xue M, He M, Huang ZJ, Scanziani M (2013) Inhibition of inhibition in visual cortex: the logic of connections between molecularly distinct interneurons. Nat Neurosci 16:1068–1076.

Pho GN, Goard MJ, Woodson J, Crawford B, Sur M (2018) Task-dependent representations of stimulus and choice in mouse parietal cortex. Nat Commun 9:2596.

Pinto L, Dan Y (2015) Cell-Type-Specific Activity in Prefrontal Cortex during Goal-Directed Behavior. Neuron 87:437–450.

Pnevmatikakis EA, Soudry D, Gao Y, Machado TA, Merel J, Pfau D, Reardon T, Mu Y, Lacefield C, Yang W, Ahrens M, Bruno R, Jessell TM, Peterka DS, Yuste R, Paninski L (2016) Simultaneous Denoising, Deconvolution, and Demixing of Calcium Imaging Data. Neuron 89:285–299.

Poort J, Khan AG, Pachitariu M, Nemri A, Orsolic I, Krupic J, Bauza M, Sahani M, Keller GB, Mrsic-Flogel TD, Hofer SB (2015) Learning Enhances Sensory and Multiple Non-sensory Representations in Primary Visual Cortex. Neuron 86:1478–1490.

Raposo D, Kaufman MT, Churchland AK (2014) A category-free neural population supports evolving demands during decision-making. Nat Neurosci 17:1784–1792.

Ringach DL, Mineault PJ, Tring E, Olivas ND, Garcia-Junco-Clemente P, Trachtenberg JT (2016) Spatial clustering of tuning in mouse primary visual cortex. Nat Commun 7:12270.

Rudy B, Fishell G, Lee S, Hjerling-Leffler J (2011) Three groups of interneurons account for nearly 100% of neocortical GABAergic neurons. Dev Neurobiol 71:45–61.

Ruebel O et al. (2019) NWB:N 2.0: An Accessible Data Standard for Neurophysiology. bioRxiv.

Runyan CA, Piasini E, Panzeri S, Harvey CD (2017) Distinct timescales of population coding across cortex. Nature 548:92–96.

Runyan CA, Schummers J, Van Wart A, Kuhlman SJ, Wilson NR, Huang ZJ, Sur M (2010) Response features of parvalbumin-expressing interneurons suggest precise roles for subtypes of inhibition in visual cortex. Neuron 67:847–857.

Rustichini A, Padoa-Schioppa C (2015) A neuro-computational model of economic decisions. J Neurophysiol 114:1382–1398.

Sahara S, Yanagawa Y, O’Leary DD, Stevens CF (2012) The fraction of cortical GABAergic neurons is constant from near the start of cortical neurogenesis to adulthood. J Neurosci 32:4755–4761.

Schoups A, Vogels R, Qian N, Orban G (2001) Practising orientation identification improves orientation coding in V1 neurons. Nature 412:549–553.

Sohya K, Kameyama K, Yanagawa Y, Obata K, Tsumoto T (2007) GABAergic neurons are less selective to stimulus orientation than excitatory neurons in layer II/III of visual cortex, as revealed by in vivo functional Ca2+ imaging in transgenic mice. J Neurosci 27:2145–2149.

Song YH, Kim JH, Jeong HW, Choi I, Jeong D, Kim K, Lee SH (2017) A Neural Circuit for Auditory Dominance over Visual Perception. Neuron 93:940–954 e946.

Taniguchi H, He M, Wu P, Kim S, Paik R, Sugino K, Kvitsiani D, Fu Y, Lu J, Lin Y, Miyoshi G, Shima Y, Fishell G, Nelson SB, Huang ZJ (2011) A resource of Cre driver lines for genetic targeting of GABAergic neurons in cerebral cortex. Neuron 71:995–1013.

Tanimoto N, Sothilingam V, Kondo M, Biel M, Humphries P, Seeliger MW (2015) Electroretinographic assessment of rod- and cone-mediated bipolar cell pathways using flicker stimuli in mice. Sci Rep 5:10731.

Teeters JL et al. (2015) Neurodata Without Borders: Creating a Common Data Format for Neurophysiology. Neuron 88:629–634.

Thomson AM, Lamy C (2007) Functional maps of neocortical local circuitry. Front Neurosci 1:19–42.

Viswanathan P, Nieder A (2015) Differential impact of behavioral relevance on quantity coding in primate frontal and parietal neurons. Curr Biol 25:1259–1269.

Vogelstein JT, Packer AM, Machado TA, Sippy T, Babadi B, Yuste R, Paninski L (2010) Fast nonnegative deconvolution for spike train inference from population calcium imaging. J Neurophysiol 104:3691–3704.

Wang XJ (2002) Probabilistic decision making by slow reverberation in cortical circuits. Neuron 36:955–968.

Wang XJ, Yang GR (2018) A disinhibitory circuit motif and flexible information routing in the brain. Curr Opin Neurobiol 49:75–83.

Wang XJ, Tegner J, Constantinidis C, Goldman-Rakic PS (2004) Division of labor among distinct subtypes of inhibitory neurons in a cortical microcircuit of working memory. Proc Natl Acad Sci U S A 101:1368–1373.

Wong KF, Wang XJ (2006) A recurrent network mechanism of time integration in perceptual decisions. J Neurosci 26:1314–1328.

Yoshimura Y, Callaway EM (2005) Fine-scale specificity of cortical networks depends on inhibitory cell type and connectivity. Nat Neurosci 8:1552–1559.

Yoshimura Y, Dantzker JL, Callaway EM (2005) Excitatory cortical neurons form fine-scale functional networks. Nature 433:868–873.

Zhong L, Zhang Y, Duan CA, Pan J, Xu N-l (2018) Dynamic and causal contribution of parietal circuits to perceptual decisions during category learning. bioRxiv.

Znamenskiy P, Kim M-H, Muir DR, Iacaruso MF, Hofer SB, Mrsic-Flogel TD (2018) Functional selectivity and specific connectivity of inhibitory neurons in primary visual cortex. bioRxiv.

